# Defective BRCA1-mediated DNA end resection drives tandem duplication formation and *FANCM* synthetic lethality

**DOI:** 10.64898/2026.02.20.706968

**Authors:** Namrata M. Nilavar, Alberto Marin-Gonzalez, Francesca Menghi, Daniel Nguyen, Nicholas A. Willis, Ellen Wientjens, Bing Xia, Jos Jonkers, Edison T. Liu, Ralph Scully

## Abstract

*BRCA1*-linked cancer genomes contain abundant genome-wide ∼10 kb ‘Group 1’ tandem duplications (TDs) that are drivers of tumorigenesis. Group 1 TD formation is recapitulated at a chromosomal Tus/*Ter* site-specific replication fork barrier in DNA end resection-defective mouse embryonic stem (mES) cells lacking *Brca1* exon 11. To explore relationships between DNA end resection and Group 1 TD formation, we analyzed *Brca1* coiled coil (CC) domain mutants—separation-of-function alleles that are impaired for homologous recombination but competent for DNA end resection. Notably, *Brca1* CC mutants retain the ability to suppress Group 1 TDs in the Tus/*Ter* system and in a mouse model of *Brca1*-linked tumorigenesis. These data show that *Brca1* CC domain mutant cancers follow a path of tumorigenesis distinct from that of other pathogenic *Brca1* alleles. FANCM is a TD co-suppressor, the loss of which is synthetic lethal/sick in combination with *Brca1* exon 11 mutation. In contrast, *Fancm* deletion is well-tolerated by *Brca1* CC mutant mES cells. Thus, Group 1 TD formation and *Fancm* synthetic lethality are linked phenotypes related to defective BRCA1-mediated DNA end resection.

## Introduction

The hereditary breast/ovarian cancer (HBOC) predisposition genes *BRCA1* and *BRCA2* regulate homologous recombination (HR), a DNA repair pathway involved in the error-free repair of double strand breaks (DSBs), protection of replication forks and the prevention of genomic instability ^1,2^. Several features of *BRCA1*- and *BRCA2*-linked cancers are shared, reflecting their roles as key HR genes. These features include elevated levels of genomic instability, a specific mutational signature (SBS3), and sensitivity to inhibitors of poly(ADP-ribose) polymerase (PARP) ^3,4^. However, the genomes of *BRCA1*-linked breast and ovarian cancers uniquely reveal abundant ∼10 kb (‘Group 1’) non-homologous tandem duplications (TDs), distributed widely across the genome ^3,5^. ‘Non-homologous’ here refers to the absence of sequence homology between the two TD breakpoints, which define the edges of the duplicated segment and are apposed to form the non-homologous TD breakpoint junction^6^. *BRCA1*-linked Group 1 TDs differ from the more common types of cancer-related TDs, which are typically >100 kb in size. Strong (∼95%) mutual concordance between *BRCA1* loss and Group 1 ‘TD phenotype’ (TDP) in human breast and ovarian cancer suggests a causal relationship between *BRCA1* loss and Group 1 TD formation ^7^. Indeed, conditional inactivation of *Brca1* in the mouse mammary epithelium recapitulates the Group 1 TD phenotype ^8^. Loss of *BRCA2* is not associated with Group 1 TDP in either human or mouse tumorigenesis, nor is *Brca2* required for the formation of the Group 1 TD in the mouse cancer model. These observations raise several questions: what mechanisms normally suppress Group 1 TDs, and what functions specific to BRCA1 mediate Group 1 TD suppression?

BRCA1 and BRCA2 operate at distinct steps of the HR pathway ^1,2,9^. The BRCA1-BARD1 heterodimer promotes short-range DNA end resection by MRE11/RAD50/NBS1 through interactions with the scaffolding protein, CtIP ^10–12^. BRCA1-BARD1 also activates long-range resection mediated by EXO1 or by the Bloom’s (BLM) or Werner’s (WRN) syndrome helicases in association with DNA2 ^13,14^. These resection activities convert the blunt or near-blunt ends of a DSB to single stranded (ss)DNA with long 3’ tails, which are then rapidly bound by the ssDNA binding protein complex RPA. BRCA2 displaces RPA from ssDNA and loads RAD51 in its place, forming the RAD51 nucleoprotein filament, which then executes the search for a homologous template ^15–18^. In addition to its role in DNA end resection, BRCA1-BARD1 interacts with the product of another HBOC gene, *PALB2*, facilitating BRCA2 recruitment and RAD51 loading ^19–23^. BRCA1 also acts at later stages of HR, facilitating strand exchange with a homologous template by the RAD51 nucleoprotein filament^24^. BRCA1 binds PALB2 directly *via* its coiled coil (CC) domain, mutations in which disrupt the interaction^21,22^. Consequently, BRCA1 CC domain mutants are defective for HR but competent for DNA end resection. Human germ-line *BRCA1* CC missense alleles are exceptionally rare and these alleles are classified as variants of uncertain significance (VUS). However, a coiled coil missense mutation in the mouse *Brca1* CC domain, p.L1363P, was found to promote mouse mammary tumorigenesis, suggesting that its human counterpart p.L1407P, a patient-derived variant, is pathogenic ^25^. Consistent with the separation-of-function properties of BRCA1 CC domain mutants, mice that are compound heterozygotes of *Brca1* Δ11 and *Brca1* ΔCC display interallelic complementation in mouse development ^26^. BRCA1’s function in DNA end resection may explain its role in single strand annealing (SSA) – a BRCA2/RAD51-independent DSB repair mechanism whereby two neighboring homologous repeats either side of a DSB can be collapsed to one copy by annealing of the complementary sequences exposed by DNA end resection ^1,2^.

To study mechanisms of stalled fork repair in mammalian cells, we previously adapted the *Escherichia coli* Tus/*Ter* replication fork barrier (RFB) to provoke bidirectional site-specific replication fork stalling on a mammalian chromosome ^27^. By targeting an HR reporter containing an array of six 23 bp *Ter* sequences to a defined chromosomal locus, we were able to quantify HR and other repair outcomes triggered by fork stalling at the Tus/*Ter* RFB. We found that conservative HR by short tract gene conversion (STGC) at Tus/*Ter* is mediated by the canonical BRCA1-BRCA2-Rad51 pathway, whereas replicative long tract gene conversion (LTGC) at Tus/*Ter* is independent of BRCA1 and BRCA2 ^27^. In addition, non-homologous Group 1 TDs form at Tus/*Ter* in *Brca1* mutant mouse embryonic stem (mES) cells, but not in *Brca2* mutants ^7^. This pattern of genetic dependency for TD formation suggests that the Tus/*Ter* system recapitulates the *BRCA1*-linked Group 1 TD mechanism observed in cancer.

Analysis of Tus/*Ter*-induced Group 1 TD formation in mES cells identified BRCA1, BARD1 and CtIP as major TD suppressors in wild type (WT) cells ^7^. The CtIP connection suggested a possible role for DNA end resection in TD suppression. Notably, Tus/*Ter*-induced TDs in DNA end resection-defective *Brca1* mutants lacking the large central in-frame exon 11 (*Brca1*^Δ11^) are elevated ∼15-fold by loss of either of two stalled fork motor proteins, FANCM or BLM, whereas loss of FANCM or BLM has minimal impact on TD formation in cells expressing WT *Brca1* ^7,28^. We found that FANCM and BLM act in concert as TD suppressors and that loss of *Fancm* is synthetic lethal/synthetic sick with *Brca1*^Δ11^ mutation. Indeed, specific inactivation of FANCM ATPase function is sufficient to disable *Brca1*^Δ11/–^ cell growth ^28^. A whole-genome CRISPR/Cas9 screen in primary human cells also identified a specific synthetic lethal relationship between *BRCA1* deletion and *FANCM* loss, but not between *BRCA2* deletion and *FANCM* loss ^29^. These findings, if verified in cancer models, identify FANCM as a potential target for therapy in *BRCA1*-linked cancer. We proposed that TDs form by a replication restart/replication bypass mechanism, in which FANCM/BLM suppresses fork restart at the Tus/*Ter* RFB, while BRCA1 promotes SSA-mediated collapse of a nascent TD back to single copy ^1,6^. Such a mechanism could explain how combined inactivation of FANCM/BLM and BRCA1-mediated DNA end resection disable different steps of the TD suppression mechanism and thereby synergize to induce very high levels of TDs and, potentially, other genomic instability outcomes^28,30^.

However, the explicit relationships between BRCA1-mediated DNA end resection, Group 1 TD formation and *FANCM* synthetic lethality remain to be rigorously tested.

Here, we study these relationships by analyzing the stalled fork repair functions of *Brca1* CC domain mutants—separation-of-function mutants that disrupt HR without affecting BRCA1-mediated DNA end resection. Our findings suggest that Group 1 TD formation and *Fancm* synthetic lethality are tightly linked phenotypes arising from defective BRCA1-mediated DNA end resection.

## Results

### A haploid system for genetic analysis of *Brca1* in stalled fork repair

To support a genetic analysis of *Brca1* in stalled fork repair, we generated a system that enables targeted mutagenesis of *Brca1* as a haploid genetic element in mES cells. This approach guarantees that subsequent CRISPR/Cas9-mediated mutagenesis of *Brca1* cannot induce unplanned changes in a second *Brca1* allele that might produce unpredictable alterations in *Brca1* function ^28,31^. We first determined whether *Brca1*^+/–^ mES cells display an overt haploinsufficiency phenotype in DSB repair or stalled fork repair. We targeted a previously described *Ter*-HR reporter^27^ to the *Rosa26* locus of *Brca1*^fl5–13/+^ cells, in which *Brca1* exons 5-13 are flanked by *loxP* sites (**Figure 1A**, **Supplemental Figure S1A**) ^32^. The *Brca1*^Δ5–13^ allele is functionally null. We then generated Cre-treated derivatives in which *Brca1* is either haploid (*Brca1*^Δ5–13/+^) or diploid (*Brca1*^fl5–13/+^) (**Supplemental Figure S1B**). In assays of DSB-induced HR or Tus/*Ter*-induced HR, *Brca1*^fl5–13/+^ and *Brca1*^Δ5–13/+^ HR reporter cells were indistinguishable (**Supplemental Figure S1C** and **S1D**). Thus, *Brca1* is not overtly haploinsufficient for DSB-induced or stalled fork HR in mES cells.

**Figure 1.**
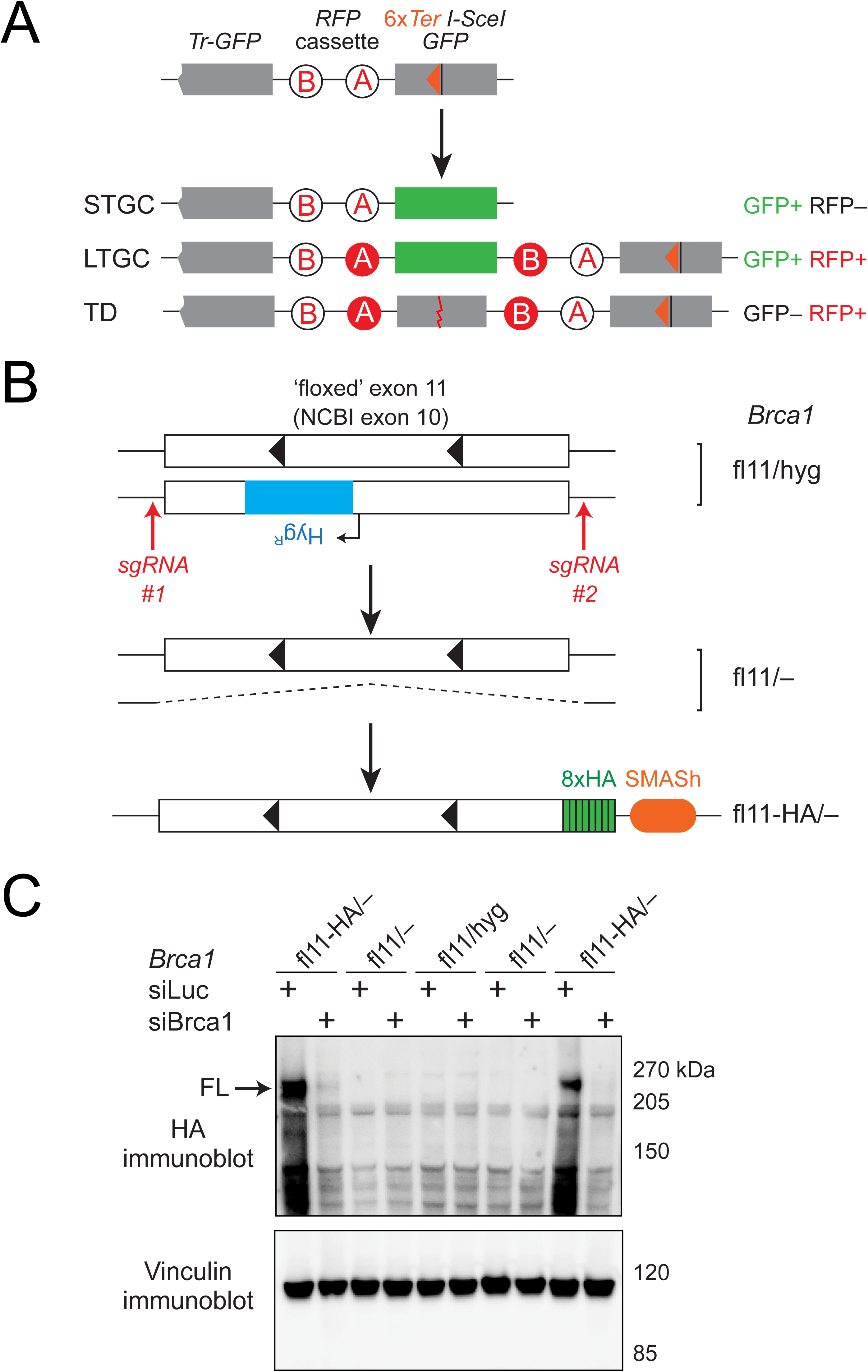
A haploid system for genetic analysis of *Brca1* in stalled fork repair. **A.** Schematic of *Ter*-HR reporter for quantitation of Tus/*Ter*-induced and DSB-induced HR. *Tr-GFP*: 5’ truncated *GFP* allele acts as donor during HR. *RFP* cassette elements A and B: artificial exons encoding WT *RFP*, placed in unproductive orientation. Promoters and polyadenylation signals not shown. STGC: short tract gene conversion. LTGC: long tract gene conversion. TD: Group 1 tandem duplication. Each repair product is associated with fluorescent products as shown. Red line in TD product: non-homologous TD breakpoint junction. **B.** Outline of method for generating haploid mES cells expressing *Brca1*-HA. Black triangles: *LoxP* sites flanking *Brca1* exon 11 (NCBI exon 10). Deleted *Brca1* allele is not shown in cartoon of *Brca1*^fl11-HA/–^ allele. **C.** Identification of HA-tagged gene product in *Brca1*^fl11-HA/–^ cells. Upper panel: anti-HA immunoblotting of whole cell extracts. Note abolition of BRCA1-HA signal by *Brca1* siRNA but not by control *Luciferase* siRNA (siLuc). FL: full length *Brca1* gene product. Lower panel: Vinculin loading control.

We next engineered haploid derivatives of *Ter*-HR reporter mES cells that carry one conditional allele of *Brca1*, *Brca1*^fl11^, in which the large central in-frame exon 11, corresponding to NCBI exon 10 (RefSeq NM_009764), can be conditionally deleted using Cre-mediated recombination^33^. The Cre-deleted allele, *Brca1*^Δ11^, is a viable hypomorph that is defective for DNA end resection^34^. The second allele, *Brca1*^hyg^, contains a hygromycin resistance gene that disrupts *Brca1* gene expression (**Figure 1B**). To delete the *Brca1*^hyg^ allele, we used two sgRNAs to target Cas9 to the extreme 5’ and 3’ ends of the *Brca1* open reading frame (ORF) and screened for clones in which one full allele had been deleted (**Figure 1B**). All derived clones were *Brca1*^fl11/–^, likely reflecting inviability of *Brca1*^hyg/–^ cells. We selected a *Brca1*^fl11/–^ clone that contained no indels in the retained *Brca1*^fl11^ allele at the 5’ and 3’ sgRNA target sites. We next used CRISPR/Cas9-assisted gene targeting to introduce an in-frame linker peptide, followed by an 8x influenza hemagglutinin (HA) epitope tag sequence to the 3’ end of the *Brca1* ORF, together with a SMASh degron^35^ and a neomycin expression cassette fused to a self-cleaving peptide T2A, to generate *Brca1*^fl11-HA/–^ cells (**Figure 1B**). The SMASh degron is also self-cleaving so that, in the absence of the degron-activating drug asuprenavir, the C-terminus of BRCA1 is fused to only the linker peptide and the 8xHA epitope tag. The full-length ∼220 kDa gene product, BRCA1-8xHA, was detectable by western blotting and was abolished by siRNA-mediated depletion of *Brca1* mRNA (**Figure 1C**). Anti-HA immunoblotting also revealed lower molecular weight *Brca1* products, reflecting products of alternative splicing or of proteolytic cleavage of full-length BRCA1. We tested the HR functions of parental *Brca1*^fl11/hyg^, *Brca1*^fl11/–^ and *Brca1*^fl11-HA/–^ cells and found them to be equivalent (**Supplemental Figure 1E**).

## Generation of *Brca1* coiled coil domain mutants

To engineer *Brca1* CC domain mutants, we used CRISPR/Cas9-assisted gene targeting to introduce the missense mutation *Brca1* p.L1363P into *Brca1*^fl11-HA/–^ cells (See STAR Methods). This allele is equivalent to the human *BRCA1* p.L1407P missense variant which shows impaired PALB2 binding ^21^. Because it is a rare allele in the human population, *BRCA1* p.L1407P is classified in ClinVar as a variant of uncertain significance (VUS). However, *Brca1* p.L1363P promotes tumorigenesis in a mouse model of mammary carcinoma ^25^. We identified several independent *Brca1*^L1363P-HA/–^ clones that express the gene product at levels equivalent to WT BRCA1, as well as two non-targeted clones in which small, in-frame deletions of 9 or 12 bp had been introduced at the sgRNA target site within the CC-coding region (**Figures 2A** and **2B** and **Supplemental Figure S2A**). These deletion alleles were termed *Brca1*^CCΔ9bp-HA/–^ and *Brca1*^CCΔ12bp-HA/–^, respectively.

**Figure 2.**
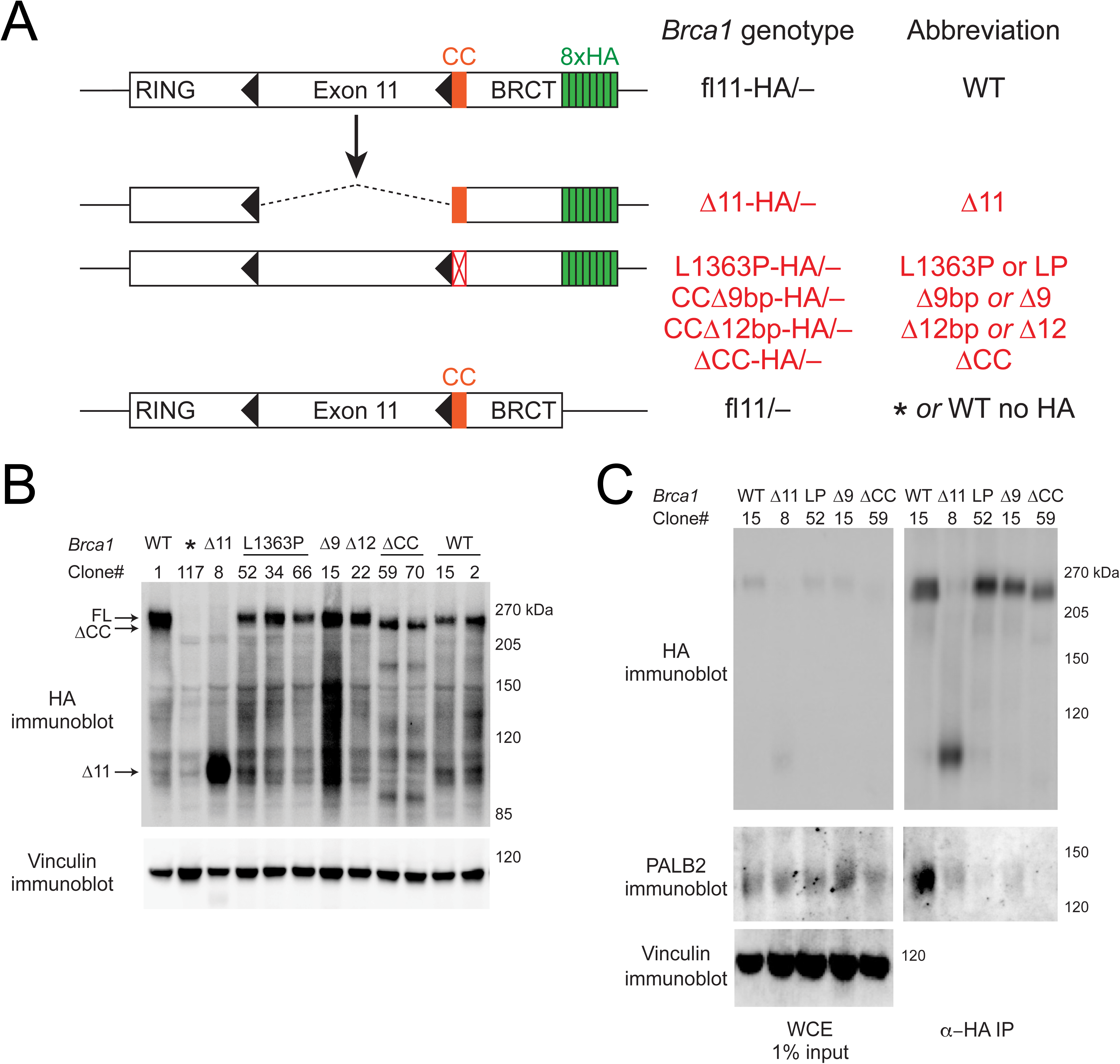
Generation of *Brca1* coiled coil domain mutants. **A.** Schematic of *Brca1* separation-of-function alleles generated by Cre-mediated recombination (*Brca1*^Δ11-HA/–^) or by CRISPR/Cas9-mediated gene editing. *Brca1* allele names are shown, together with abbreviations used in this and subsequent figures. **B.** Identification of HA-tagged gene product in *Brca1*^HA/–^cells with mutations as shown. *: parental *Brca1*^fl11/–^ clone, lacking HA-tag. Upper panel: anti-HA immunoblotting of whole cell extracts. FL: arrow shows full-length BRCA1. ΔCC: arrow shows *Brca1*^ΔCC-HA/–^ gene product. Lower panel: vinculin loading control. **C.** Detection of PALB2 association with products of different *Brca1*^HA/–^ alleles. Left panels: whole cell extracts. Right panels: anti-HA immunoprecipitates. Immunoblots were probed with the antibodies shown. Lower panel: Vinculin loading control in whole cell extracts.

We also used CRISPR/Cas9 dual incisions to generate larger deletions within the *Brca1* CC-encoding region (*Brca1*^ΔCC-HA/–^), spanning the two exons (NCBI exons 11 and 12) that encode this domain. We retrieved two clones, one of which (#70) contained a predicted in-frame deletion and one (#59) a frame-shifting deletion (**Supplemental Figure S2B**). Despite these genetic differences, the two alleles encoded HA-tagged *Brca1* gene products of equal size that were slightly smaller than the WT BRCA1 protein (**Figure 2B**). Analysis of *Brca1* mRNA by RT-qPCR across specific exon-exon junctions and by direct sequencing of *Brca1* cDNA revealed that NCBI exons 11 and 12 were skipped in both ΔCC clones, resulting in an in-frame deletion of 81 amino acids, effectively deleting the entire coiled coil domain (**Supplemental Figures S2C-S2E**). This exon-skipping explains why ΔCC allele #59 does not truncate the protein, and why the two different ΔCC alleles encode similar gene products. We presume that the emergence of this exon-skipping adaptation is achieved through selection against cells that fail to skip NCBI exons 11 and 12, since the exon-skipped mRNA is virtually undetectable in WT cells (**Supplemental Figure S2C**). It is not clear how reconfigurations of the splicing apparatus enable the observed exon skipping to occur.

We used Cre-mediated recombination to delete exon 11 in *Brca1*^fl11-HA/–^ cells (see STAR Methods) and identified the *Brca1*^Δ11-HA/–^ gene product by western blotting (**Figures 2A** and **2B**). We used immunostaining and immunofluorescence microscopy to assess the subcellular localization of each *Brca1* gene product (**Supplemental Figure 3**). Consistent with previous reports, BRCA1Δ11-HA and all other BRCA1 proteins localized to characteristic nuclear foci ^36,37^. The anti-HA signal of WT and CC mutant proteins co-localized with foci detected by an antibody raised against a polypeptide encoded by *Brca1* exon 11, whereas this exon 11-specific BRCA1 signal was absent in *Brca1*^Δ11-HA/–^ cells (**Supplemental Figure 3**). Similarly, the anti-HA signal was absent in *Brca1*^fl11/–^ cells, which lack the HA epitope tag.

We next assessed PALB2 binding by BRCA1 in each genetic background, using anti-HA immunoprecipitation (IP) followed by immunoblotting for endogenous PALB2 (**Figure 2C**). Despite equivalent levels of PALB2 in cell extracts, IPs from BRCA1 CC mutants revealed a severely attenuated PALB2 signal, with BRCA1ΔCC-HA revealing the most severe PALB2 binding defect. Interestingly, BRCA1Δ11-HA also revealed reduced binding to PALB2 in comparison to WT BRCA1-HA, as was noted previously^26^.

## Defective HR and intact DNA end resection in *Brca1* coiled coil mutants

*Brca1* CC mutants are known to be defective for DSB-induced HR, but their function in stalled fork HR has not yet been studied. We showed previously that *Brca1*^Δ11/hyg^ cells exhibit reduced STGC in response to a Tus/*Ter* RFB, while Tus/*Ter*-induced LTGC is independent of *Brca1* and *Brca2* ^27^. To study stalled fork HR in *Brca1* CC mutant cells, we quantified Tus/*Ter*-induced and I-SceI-induced HR in *Brca1*^HA/–^ mutant clones *vs*. isogenic WT controls, which had been exposed to the same CRISPR/Cas9 or Cre treatments but had retained WT *Brca1*. Mean values of Tus/*Ter*-induced STGC (averaged across the clones shown) were 64 % of WT in *Brca1*^Δ11-HA/–^cells, 23 % in *Brca1*^ΔCC-HA/–^ cells and 37 % in *Brca1*^L1363P-HA/–^ cells (**Figures 3A**-**3C**). Equivalent mean values of I-SceI-induced STGC were 36 % of WT in *Brca1*^Δ11-HA/–^ cells, 27 % in *Brca1*^ΔCC-HA/–^ cells and 50 % in *Brca1*^L1363P-HA/–^ cells (**Supplemental Figure S4A**-**4C**). STGC defects in the *Brca1*^CCΔ9bp-HA/–^ clone resembled those of *Brca1*^L1363P-HA/–^ clones. There was no consistent impact of *Brca1* CC domain mutation on Tus/*Ter*-induced LTGC (**Figures 3B**-**3C**).

**Figure 3.**
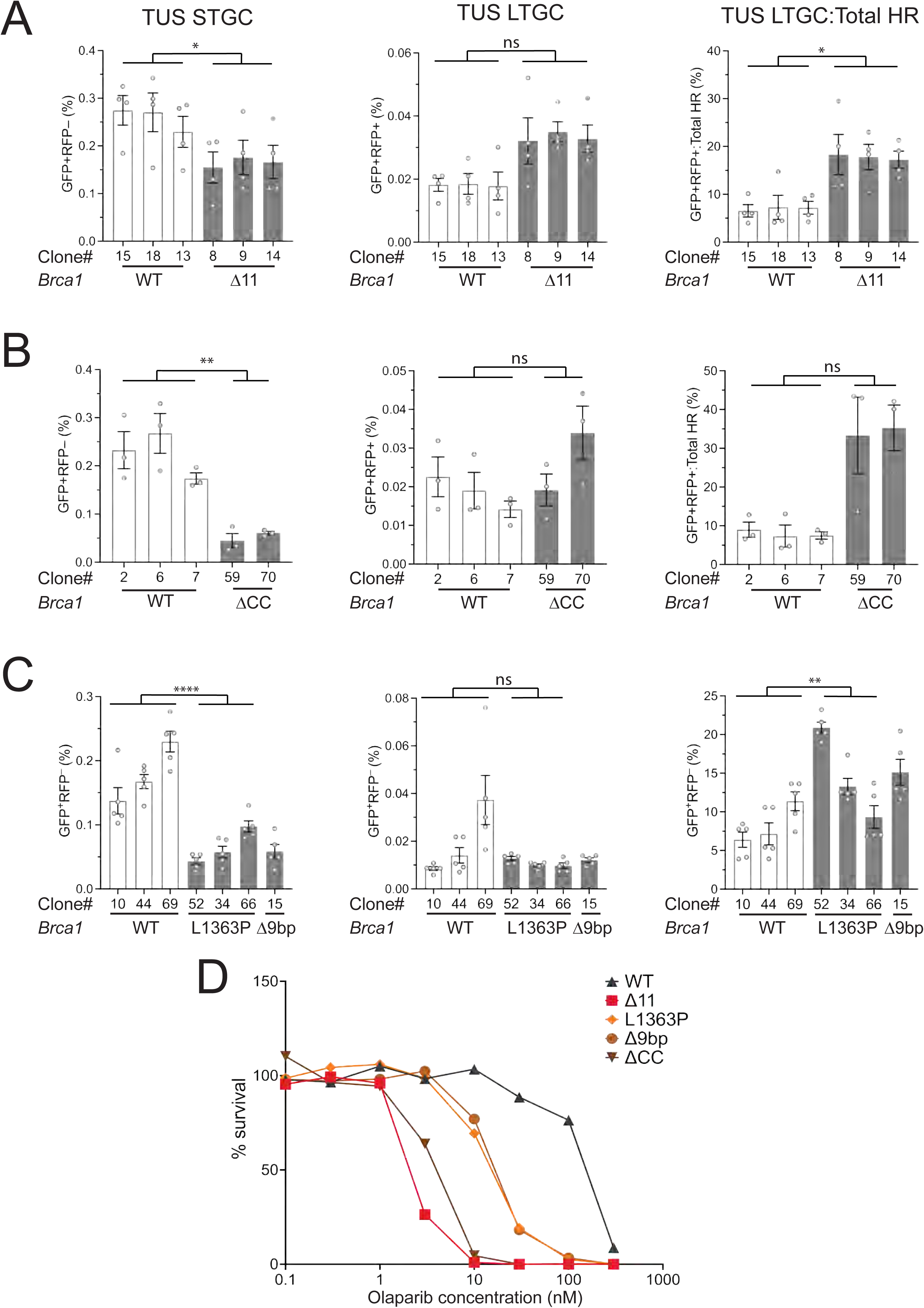
Tus/*Ter*-induced HR and PARPi sensitivity of *Brca1* coiled coil mutants. HR was quantified in independent *Brca1* mutant (grey) or isogenic *Brca1* WT (white) clones as described in STAR Methods. *Brca1* allele abbreviations as in Figure 2A. **A.** *Brca1*^Δ11-HA/–^ (n=4). **B.** *Brca1*^ΔCC-HA/–^ cells (n=3). **C.** *Brca1*^L1363P-HA/–^ and *Brca1*^CCΔ9bp-HA/–^ (n=5). Data shows mean and SEM of independent experiments. Statistical analysis: RM-one way ANOVA with Geisser-Greenhouse correction: ns: not significant, * p<0.05, ** p<0.005, *** p<0.0005, ****p<0.0001. **D.** Colony formation of different *Brca1*^HA/–^ genotypes in response to concentrations of Olaparib as shown (See STAR Methods for details). Data shows mean of 5 independent biological replicates normalized to colony numbers in untreated samples of the same genotype (n=5).

To further characterize the HR functions of the different *Brca1* mutant clones, we assessed sensitivity to the PARP inhibitor Olaparib by colony formation assay (**Figure 3D**). IC50 values were ∼150 nM for WT cells, 2 nM for *Brca1*^Δ11-HA/–^ cells, 3.5 nM for *Brca1*^ΔCC-HA/–^ cells, and ∼15 nM for both *Brca1*^L1363P-HA/–^ and *Brca1*^CCΔ9bp-HA/–^ clones. Notably, *Brca1* Δ11 mutants were among the most sensitive to Olaparib and the most impaired in I-SceI-induced STGC, but revealed the mildest defect in Tus/*Ter*-induced STGC. This discordance suggests that an individual *Brca1* mutant’s PARPi sensitivity correlates better with its function in DSB-induced HR than in stalled fork HR.

To determine the DNA end resection phenotypes of *Brca1* exon 11-deleted and CC domain mutants, we quantified RPA accumulation at a defined chromosomal DSB site using chromatin-immunoprecipitation-sequencing (ChIP-seq; see STAR Methods) (**Figure 4A**). Three hours after electroporation of CRISPR/Cas9 ribonucleoprotein (RNP) complexes targeting the *I-SceI* site within the *Ter*-HR reporter, we observed strong RPA enrichment in regions flanking the DSB site in *Brca1*^fl11-HA/–^ cells (**Figure 4B**). In contrast, the RPA ChIP-seq signal was greatly diminished in *Brca1*^Δ11-HA/–^ cells, consistent with the expected defect in DNA end resection. Notably, *Brca1* CC domain mutants *Brca1*^L1363P-HA/–^, *Brca1*^Δ9bp-HA/–^ and *Brca1*^ΔCC-HA/–^ each revealed levels of RPA accumulation similar to those of *Brca1*^fl11-HA/–^ cells (**Figures 4B** and **4C**). Thus, each of the *Brca1* CC domain mutants studied here supports WT levels of DNA end resection.

**Figure 4.**
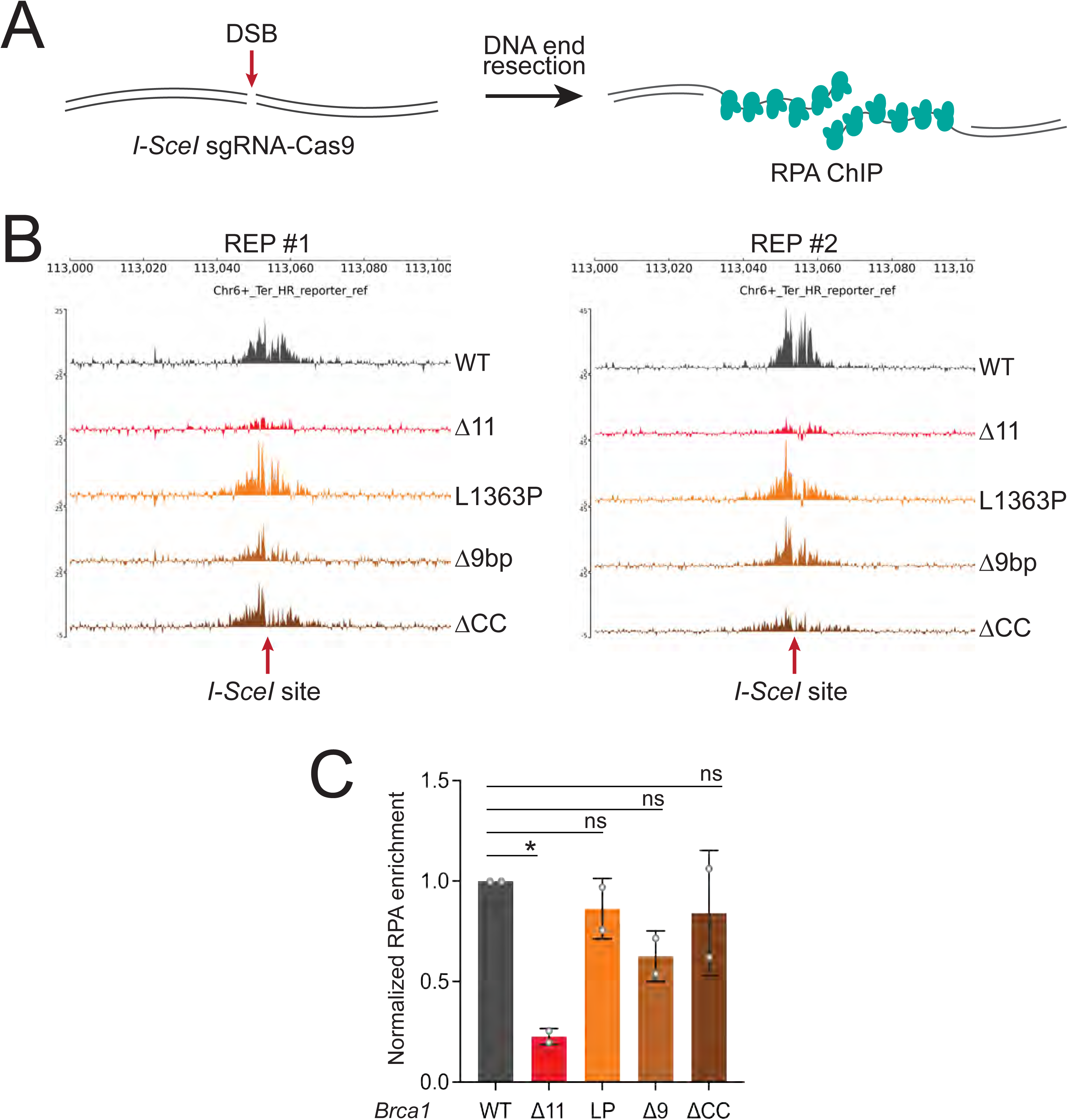
*Brca1* coiled coil mutants are competent for DNA end resection. **A.** Schematic of RPA accumulation on ssDNA in response to DNA end resection of a Cas9 induced DSB, targeted to the *I-SceI* site within the HR reporter at *Rosa26*. Intensity of RPA ChIP-seq signal reflects the efficiency of DNA end resection. **B.** Cas9 DSB-induced RPA ChIP-seq signals in two independent experiments (REP#1 and REP#2) using *Brca1* genotypes shown. *Brca1* allele abbreviations as in Figure 2A. See STAR Methods for details. **C**. Quantitation of RPA enrichment averaged over 30 kb window flanking *I-SceI* site. Signals were normalized to WT for each of the replicates. Statistical analysis: unpaired *t*-test with Welch’s correction. ns: not significant, p *<0.05.

## *Brca1* CC domain mutants are competent for suppression of Tus/*Ter*-induced TDs

We showed previously that FANCM depletion strongly enhances Tus/*Ter*-induced TD formation in *Brca1*^Δ11/hyg^ cells ^7,28^. To test whether *Brca1* CC domain mutants are competent for TD suppression, we quantified Tus/*Ter*-induced TD frequencies in cultures of *Brca1*^fl11-HA/–^, *Brca1*^L1363P-HA/–^, *Brca1*^CCΔ9bp-HA/–^, *Brca1*^ΔCC-HA/–^ and *Brca1*^Δ11-HA/–^ cells that had been depleted of FANCM using siRNA. Controls in this experiment included non-targeting luciferase siRNA (siLuc, negative control) and combined siRNA directed to *Brca1* and *Fancm* (positive control). The former provides the baseline level of Tus/*Ter*-induced TDs in each clone; the latter tests whether each clone can, in principle, form TDs when BRCA1 and FANCM are co-depleted.

Indeed, each cell line revealed induction of Tus/*Ter*-induced GFP^−^RFP^+^ products (i.e., TDs) following siRNA-mediated co-depletion of BRCA1 and FANCM (**Figures 5A** and **5B**). When depleted of FANCM alone, *Brca1*^fl11-HA/–^ cells revealed minimal TD induction, whereas *Brca1*^Δ11-HA/–^ cells revealed high levels of Tus/*Ter*-induced TDs, as expected. Strikingly, unlike in *Brca1*^Δ11-HA/–^ cells, FANCM depletion failed to stimulate Tus/*Ter*-induced TDs in *Brca1*^L1363P-HA/–^or *Brca1*^CCΔ9bp-HA/–^ cells, with TD frequencies equivalent to those observed in WT *Brca1*^fl11-HA/–^cells (**Figure 5A**). *Brca1*^ΔCC-HA/–^ cells revealed a modest increase in Tus/*Ter*-induced TDs, but significantly less than in *Brca1*^Δ11-HA/–^ cells (**Figure 5B**). Thus, *Brca1* CC domain mutants retain the ability to suppress Group 1 TD formation at a Tus/*Ter* RFB. We showed previously that I-SceI-induced GFP^−^RFP^+^ products are not non-homologous TDs, but represent a rare, aberrant outcome of sister chromatid recombination ^38^. *Fancm* depletion in the context of I-SceI-induced repair failed to generate a significant increase in GFP^−^RFP^+^ products in any of the *Brca1* mutant clones studied (**Supplemental Figures S4D**-**S4F**). In summary, the data suggest that the critical determinant of BRCA1-mediated Group 1 TD suppression at a Tus/*Ter* RFB is DNA end resection function, but not overall HR capacity.

**Figure 5.**
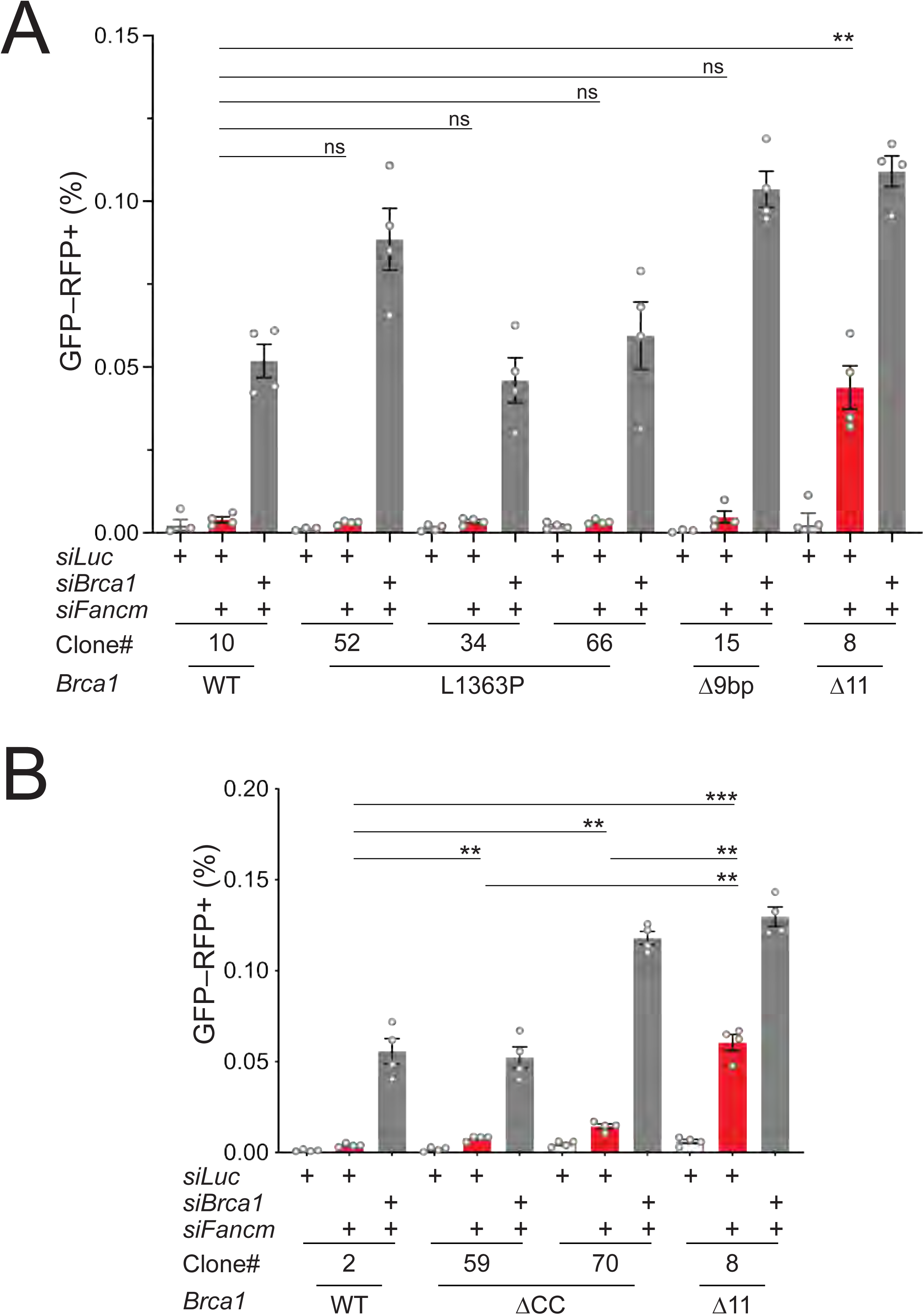
*Brca1* coiled coil mutants suppress Tus/*Ter*-induced tandem duplications. Tus/*Ter*-induced GFP^−^RFP^+^ repair products in *Brca1* WT and mutant genotypes, in the presence of siRNAs against *Fancm* and *Brca1*. *Brca1* allele abbreviations as in Figure 2A. See STAR Methods for details. siRNA against luciferase (si*Luc*): non-specific siRNA negative control. Gray columns: combined si*Fancm* and si*Brca1* induces TDs in all cell lines (positive control). Red columns: si*Fancm* and si*Luc* test TD suppression in the *Brca1* genotypes shown. **A.** *Brca1*^L1363P-HA/–^ and *Brca1*^CCΔ9bp-HA/–^ mutants are compared with isogenic WT *Brca1*^fl11-HA/–^ cells and with *Brca1*^Δ11-HA/–^ cells (n=4). Note induction of TDs in *Brca1*^Δ11-HA/–^ cells but not in CC mutants. **B.** *Brca1*^ΔCC-HA/–^ mutants are compared with isogenic WT cells and *Brca1*^Δ11-HA/–^ cells (n=4). Statistical analysis: Unpaired *t*-test with Welch’s correction. ns: not significant, p ** < 0.005. *** < 0.0005.

### *Brca1* p.L1363P mouse mammary tumors do not display a Group 1 TD phenotype

Our findings that *Brca1* CC domain mutants are competent for suppression of Tus/*Ter*-induced TDs raised the question of whether cancers driven by *Brca1* CC mutation will or will not accumulate Group 1 TDs genome-wide during tumorigenesis and whether the *Brca1* CC mutant cancer genome will register as Group 1 TDP. While the Tus/*Ter* system is an indication of TD formation at one specific site, analysis of TDs in the *Brca1* CC mutant cancer genome would enable us to further test of the predictive power of the Tus/*Ter* system. We were unable to obtain tissue samples from human *BRCA1* CC mutant cancers, reflecting their rarity in the human population. We therefore analyzed a well-established mouse model of mammary tumorigenesis, in which *K14-Cre* or *WAP-Cre* expression in the mammary epithelium drives biallelic deletion of *Trp53* by converting *Trp53*^fl2-10/fl2-10^ to null (*Trp53*^Δ2-10/Δ2-10^) together with deletion of conditional alleles of *Brca1* (*Brca1*^fl5-13/fl5-13^) or *Brca2* (*Brca2*^fl11/fl11^) ^8,32,39^. Pulver *et al*. developed this model further, using the *Brca1* p.L1363P allele as a cancer driver ^25^. We analyzed TDs in the genomes of mammary tumors arising from *Cre*-expressing *Trp53*^fl2-10/fl2-10^ mice in three different *Brca* genotypes: first, *Brca1*^fl5-13/fl5-13^, in which Cre converts both *Trp53* and *Brca1* to null (*Brca1*^Δ5-13/Δ5-13^;*Trp53*^Δ2-10/Δ2-10^, hereafter termed ‘B1P’); second, *Brca2*^fl11/fl11^, in which Cre converts *Trp53* and *Brca2* to null (*Brca2*^Δ11/Δ11^;*Trp53*^Δ2-10/Δ2-10^, hereafter ‘B2P’); third, *Brca1*^fl5-13/L1363P^, in which Cre converts *Trp53* and the *Brca1*^fl5-13^ allele to null, resulting in functional expression of *Brca1*^L1363P^ alone (*Brca1*^Δ5-13/L1363P^*;Trp53*^Δ2-10/Δ2-10^, hereafter ‘Br1L1363P’). We used whole genome sequencing (WGS) to analyze 5 independent tumors of each genotype, using WGS of spleen tissue from each mouse to define the reference genome. We included samples from our previous analysis of Cre-driven *Brca* mutant mouse mammary tumors, including controls in which *Trp53* was the only conditional allele^8^.

To confirm loss of the floxed regions *via* Cre-mediated recombination in each tumor genome, we first manually inspected the Cre-targeted regions using the Integrative Genomic Viewer (IGV), for evidence of (i) a decrease in coverage affecting the Cre-targeted segments of the Δ alleles of *Trp53*, *Brca1* or *Brca2* (ii) a deletion rearrangement supporting the recombination of targeted regions, and/or (iii) evidence of the p.L1363P mutation (examples in **Supplemental Figure 5A**).

Br1L1363P cancers contain a single copy of the *Brca1*^Δ5-13^ allele in their genomes, and generally exhibit high stromal infiltration, as previously described^25^. Elevated stromal content can dilute tumor cell-derived genotype signals, making it challenging to determine the true extent of Cre-mediated recombination of floxed alleles in tumor cells (i.e. partial *vs*. complete). To address this potential confounder, we compared the ratio between the Δ allele frequency (DeltaFr) of the *Trp53* gene *vs*. that of the *Brca1* gene in both the Br1L1363P and B1P cancer genomes. A DeltaFr = 1 corresponds to 100% decrease in coverage across the Δ region compared to its flanking regions (i.e., no stromal contamination and complete recombination); whereas a DeltaFr = 0 corresponds to equal coverage between flanking and Δ regions (i.e., elevated stromal contamination and/or incomplete recombination). Finally, we computed the ratio between the DeltaFr relative to the *Trp53* and *Brca1* loci, working under the expectation that a complete recombination would result in *Trp53-DeltaFr:Brca1-DeltaFr* ratios of approximately 1 for the B1P and B2P models and ∼2 for the Br1L1363P models (example in **Supplemental Figure 5B**; see also STAR Methods). Indeed, this ratio was ∼1 in B1P and B2P tumors, but ∼2 in Br1L1363P tumors, reflecting the loss of one *Brca1* allele and two *Trp53* alleles, with retention of the *Brca1* p.L1363P allele (**Supplemental Figure 5C**). In one Br1L1363P tumor (T7), loss of *Brca1*^fl5-13^ was accompanied by a much larger deletion that included the flanking sequences, making calculation of this ratio unreliable. This larger deletion resulted in the loss of the entire *Brca1*^fl5-13^ allele, as reflected by the p.L1363P mutation having a variant allele frequency of ∼1 in this sample (not shown).

We assessed the TD configuration of each tumor genome on a whole genome scale. We noted abundant Group 1 TDs in 5/5 B1P tumor genomes in sufficient numbers to be scored as TDP Group 1^8^ (**Figures 6A** and **6B**). In contrast, 5/5 Br1L1363P and 5/5 B2P tumor genomes revealed no enrichment of Group 1 TDs, failed to score as TDP and revealed statistically similar distributions of TD span sizes (**Figures 6A** and **6B**). Interestingly, one B2P tumor sample, B2P-T2, showed an exceptional rearrangement pattern, with hyperabundance of TDs, mostly of span size ≤ 1 kb or > 100 kb (**Figure 6A**). However, the B2P-T2 tumor failed to score as TDP by the algorithm of Menghi *et al*. ^5^, because the regions of intense instability were not dispersed genome-wide, as is characteristic of TDP cancer genomes. Furthermore, the B2P-T2 genome revealed single nucleotide variant frequencies ∼100-fold higher than all other samples (**Supplemental Figure S5D**), suggesting a complex origin of this excessive genomic instability. Irrespective of whether the outlier tumor B2P-T2 is included or excluded from the TD span size analysis (compare **Figure 6C** with **Figure 6B**), B2P tumors and Br1L1363P tumors revealed statistically similar TD spectra mostly devoid of the *Brca1*-deficiency linked Group 1 TDs. Most TDs in B2P and Br1L1363P tumor genomes were >100 kb, as is typical of the generic cancer genome ^6,40^. Comparison to historical *Trp53*^Δ2-10/Δ2-10^ controls showed that tumor genomes from all groups in which a *Brca* gene had been targeted revealed statistically significant elevations in the abundance of the SBS3 mutation signature (**Figure 6D**) ^3^. The findings that *Brca1* p.L1363P drives accelerated tumorigenesis in the mouse^25^ and that Br1L1363P tumor genomes show the distinctive SBS3 mutation signature strongly suggest that *Brca1* p.L1363P is a pathogenic variant.

**Figure 6.**
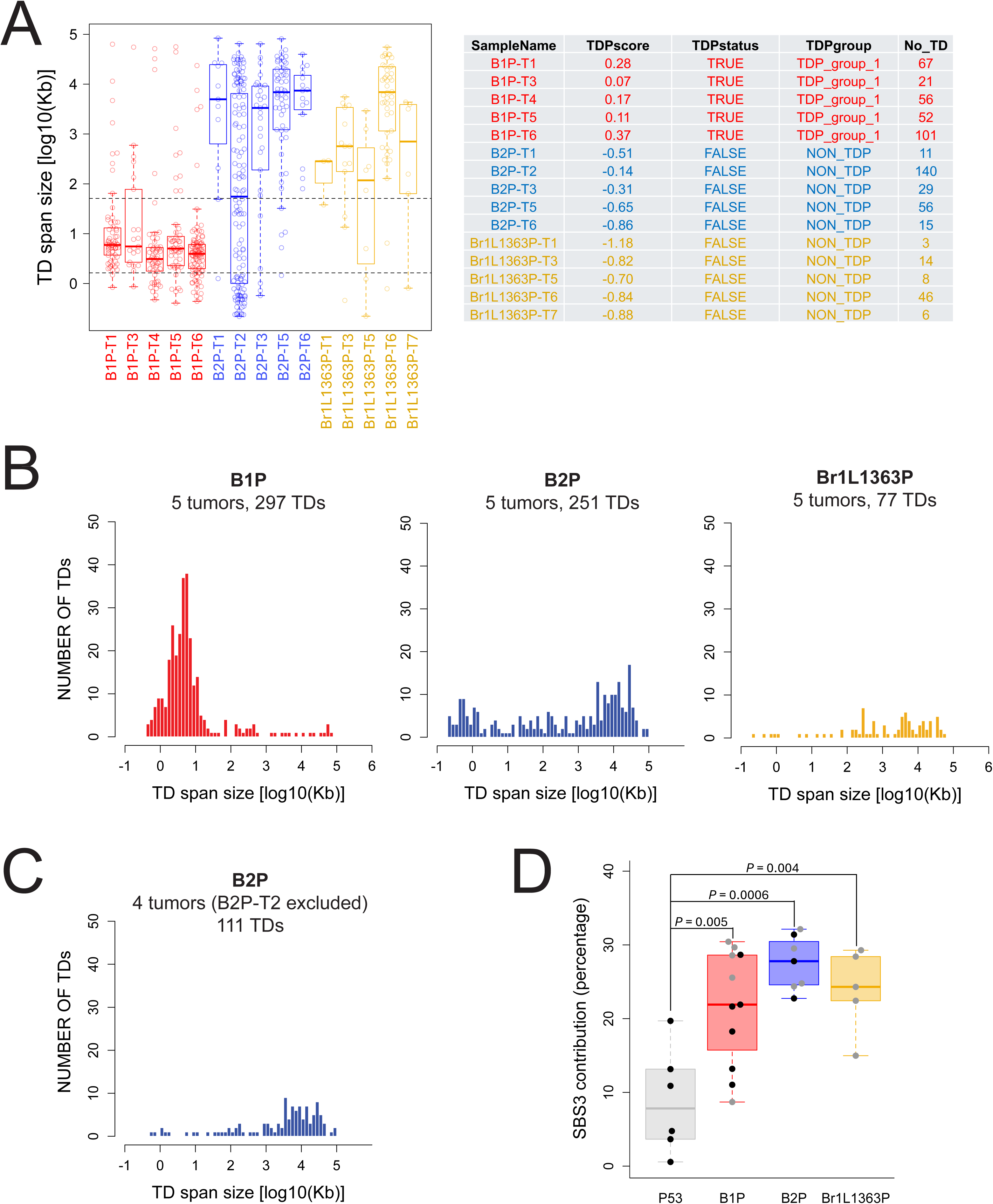
Analysis of tandem duplications in *Brca* p.L1363P mouse mammary tumors. **A.** Left panel: Distribution of TD span sizes in five mouse mammary tumor genomes of different *Brca* genotypes. B1P: *Brca1*^Δ5-13/Δ5-13^*;Trp53*^Δ2-10/Δ2-10^. B2P: *Brca2*^Δ11/Δ11^;*Trp53*^Δ2-10/Δ2-10^. Br1L1363P: *Brca1*^Δ5-13/L1363P^*;Trp53*^Δ2-10/Δ2-10^. The dotted horizontal lines denote the TD span size range for Group 1 TDs. Note abundant Group 1 TDs in B1P tumor genomes but not in B2P or Br1L1363P tumor genomes. Right panel: TDP scores for each tumor genome. 5/5 B1P tumors score as Group 1 TDP but no B2P or Br1L1363P tumors score as TDP. **B.** TD span size distribution in 5 pooled tumors of each *Brca* genotype. Mann-Whitney U test of TD size distributions: B1P vs B2P: P=1.4E-22. B1P vs Br1L1363P: P=2.8E-25. B2P vs Br1L1363P: not significant (ns). **C.** TD span size distribution in 4 pooled B2P tumors, with exclusion of outlier B2P-T2. Mann-Whitney U test of TD size distributions (with exclusion of B2P-T2): B1P vs B2P: P=1.4E-22. B2P vs Br1L1363P: ns. **D.** Quantitation of SBS3 mutation signature in the genotypes shown. This data includes both current data (grey data points) and historical data from a previous study (black data points) that includes negative control *Trp53*^Δ2-10/Δ2-10^ tumors (P53) in which *Brca* alleles are wild type. Statistical comparisons by Student’s *t*-test (two-tailed); P values as shown. B1P *vs*. B2P: ns. B1P *vs*. Br1L1363P: ns. B2P *vs*. Br1L1363P: ns.

Taken together, despite being defective for HR, *Brca1* p.L1363P remains competent for suppression of genome-wide Group 1 TD formation during tumorigenesis. The Tus/*Ter* system accurately predicted this separation-of-function phenotype in *Brca1* p.L1363P-driven tumors.

## *Brca1* CC domain mutants tolerate *Fancm* deletion

DNA end resection-defective *Brca1*^Δ11/hyg^ mES cells reveal abundant TDs when depleted of FANCM and are synthetic lethal/synthetic sick on a *Fancm* null background ^28^. To better understand the relationships between Group 1 TD formation, DNA end resection and *Fancm* synthetic lethality, we asked whether *Brca1* CC domain mutants tolerate *Fancm* deletion. To this end, we used CRISPR/Cas9-mediated dual DSB incisions to delete the DEAH (ATP binding) motif of the FANCM ATPase domain and to introduce an 85 bp frame-shift within *Fancm* exon 2, generating clones of *Fancm*^+/–^ mES cells on different *Brca* genetic backgrounds: *Brca1*^fl11-HA/–^, *Brca1*^L1363P-HA/–^, *Brca1*^Δ11-HA/–^ and *Brca2*^Δ27/Δ27^ (**Supplemental Figure S6A**). We then repeated the CRISPR/Cas9 transfection to delete the residual WT *Fancm* allele. This two-step method was necessary because the efficiency of biallelic *Fancm* deletion in *Fancm*^+/+^ cells was low. We retrieved viable *Fancm*^+/–^ or *Fancm*^−/−^ derivatives on all *Brca* genetic backgrounds. However, each *Fancm*^−/−^;*Brca1*^Δ11-HA/–^ clone was extremely slow-growing, indicating a synthetic sick phenotype. We quantified Tus/*Ter*-induced TDs in the *Fancm*^−/−^;*Brca* mutants derived above. *Fancm*^−/−^;*Brca1*^Δ11-HA/–^ clones revealed robust TD induction. In contrast, *Fancm*^−/−^;*Brca1*^L1363P-HA/–^ and *Fancm*^−/−^;*Brca2*^Δ27/Δ27^ clones revealed frequencies of Tus/*Ter*-induced TDs similar to or lower than those seen in *Fancm*^−/−^;*Brca1*^WT-HA/–^ cells (**Supplemental Figure S6B**).

We quantified the growth properties of *Fancm*^−/−^;*Brca* mutants using two independent assays: plating efficiency/colony formation assays and competitive growth assays. We noted a dramatic loss of plating efficiency in two independent clones of *Fancm*^−/−^;*Brca1*^Δ11-HA/–^ cells, in comparison to *Fancm*^+/–^;*Brca1*^Δ11-HA/–^ isogenic controls (**Figures 7A** and **7B**). In contrast, two independent clones of *Fancm*^−/−^;*Brca1*^L1363P-HA/–^ cells revealed no loss of plating efficiency in comparison to isogenic *Fancm*^+/–^;*Brca1*^L1363P-HA/–^ cells. There was a modest reduction in plating efficiency in two independent clones of *Fancm*^−/−^;*Brca2*^Δ27/Δ27^ cells ^41^. To perform competitive growth assays, we mixed ∼10% GFP^+^ *Fancm*^−/−^;*Brca1*^fl11-HA/–^ cells (WT controls) with ∼90% GFP^−^ *Fancm*^−/−^cells of the test *Brca* genotypes. We then cultivated the mixed populations over a two-week period, harvesting aliquots at defined timepoints for FACS quantitation of the GFP^+^ fraction (**Figures 7C** and **7D**). As expected, the *Brca1* WT (GFP^−^) *Fancm*^−/−^;*Brca1*^fl11-HA/–^ sample produced GFP fractions that were stable over the two-week period. In contrast, *Fancm*^−/−^;*Brca1*^Δ11-HA/–^ cultures revealed progressive increases in the GFP fraction over time, indicative of a growth defect in the GFP^−^ *Fancm*^−/−^;*Brca1*^Δ11-HA/–^ cells in comparison to the GFP^+^ *Fancm*^−/−^;*Brca1*^fl11-HA/–^ co-cultured cells. We noted a more modest competitive growth deficit in the *Fancm*^−/−^;*Brca2*^Δ27/Δ27^ cells. Surprisingly, two independent *Fancm*^−/−^;*Brca1*^L1363P-HA/–^ clones revealed a gradual *reduction* in the GFP fraction during the two-week co-culture period, indicative of a modest growth advantage over the GFP^+^ *Fancm*^−/−^;*Brca1*^fl11-HA/–^ co-cultured cells. Thus, in contrast to *Brca1* Δ11 mutants, which develop a severe growth defect following *Fancm* deletion, *Brca1* CC domain mutants are tolerant of *Fancm* loss. This result suggests that synthetic lethality/sickness following *Fancm* loss in *Brca1* mutants is correlated with defective DNA end resection function but not with overall HR capacity.

**Figure 7.**
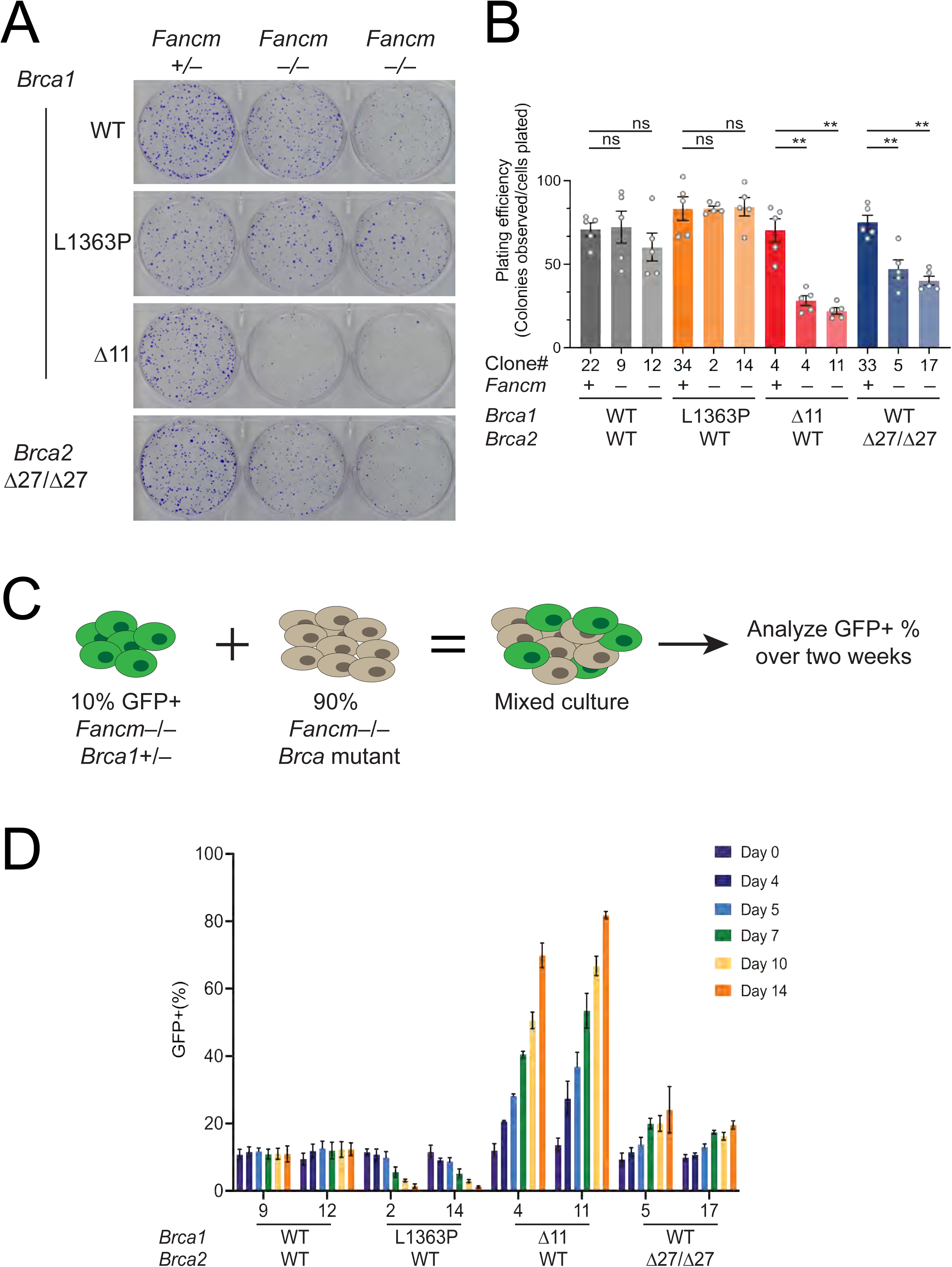
*Fancm* deletion is tolerated by *Brca1* p.L1363P cells. **A.** Representative images of colony formation in the genotypes shown. *Brca1* allele abbreviations as in Figure 2A. Note severe loss of colony forming ability of two independent *Fancm*^−/−^*Brca1*^Δ11-HA/–^ clones. **B.** Quantitation of colony formation assays in five independent biological replicates (n=5). Unpaired *t*-test with Welch’s correction. ns: not significant, p ** < 0.005. **C.** Schematic outline of competitive growth assay. **D.** Competitive growth assays of the *Fancm*^−/−^*Brca* genotypes shown. Note progressive enrichment of GFP^+^-marked *Fancm*^−/−^*Brca1*^fl11-HA/–^ cells when mixed with *Fancm*^−/−^*Brca1*^Δ11-HA/–^ cells.

## Discussion

Here we compare two classes of *Brca1* separation-of-function alleles in stalled-fork HR and in Group 1 TD formation. BRCA1 CC mutant proteins are defective for interaction with PALB2, leading to a defect in RAD51-mediated HR ^19–23^. However, unlike the majority of cancer-associated *BRCA1* mutants, BRCA1 CC mutants remain competent for DNA end resection because they retain the key BRCA1 domains (RING, tandem BRCT and exon 11-encoded region) required for this function ^13,14^. *Brca1* Δ11 and *Brca1* CC mutants are complementary separation-of-function alleles, as shown by interallelic complementation between these two alleles in mouse development^26^. A likely explanation of this phenomenon is that *Brca1* CC mutants provide DNA end resection capability, while *Brca1* Δ11 mutants facilitate PALB2 recruitment for HR. In the mES cell system studied here, we also observed reduced PALB2 binding by endogenous BRCA1 Δ11 in comparison to WT BRCA1. This defect was more severe than that noted previously in human cancer cells expressing exogenous *BRCA1* alleles^26^. Thus, although *Brca1* Δ11 mutants are DNA end resection-defective, we cannot exclude the possibility that coordination between BRCA1 Δ11 and PALB2-BRCA2-RAD51 is also impaired. The minimal PALB2-binding region of mouse BRCA1 comprises residues 1350-1377 (PDB: 7K3S).

Conceivably, the full interface between BRCA1 and PALB2 might extend to BRCA1 exon11-encoded regions neighboring the CC domain.

The effects of *Brca1* exon 11 deletion and CC mutation on stalled fork HR, DSB-induced HR and PARPi sensitivity were qualitatively similar but revealed interesting quantitative differences. *Brca1* ΔCC mutant cells, in which the entire CC domain is deleted, revealed greater sensitivity to the PARP inhibitor Olaparib and more severe STGC defects than *Brca1* L1363P or Δ9bp cells, in which the BRCA1 coiled coil domain is disrupted but not fully deleted. This difference may reflect more complete loss of PALB2 binding by BRCA1 ΔCC than by BRCA1 L1363P or BRCA1 Δ9bp. Interestingly, stalled fork STGC failed to accurately predict PARPi sensitivity.

This is most strikingly seen in *Brca1* Δ11 cells, which were among the most PARPi-sensitive but revealed the mildest defects in Tus/*Ter*-induced STGC of all *Brca1* mutants studied here.

Similarly, we previously noted that *Brca1* BRCT domain mutant mES cells reveal no defect in Tus/*Ter*-induced STGC, despite being PARPi sensitive and defective for DSB-induced STGC^27,42,43^. This discrepancy suggests that the DNA end resection function of BRCA1 is less critical for Tus/*Ter*-induced STGC than it is for I-SceI-induced STGC or for PARPi resistance. Similar effects were noted in recent studies of nickase-induced repair, where *Brca1* was found to be dispensable for DNA end resection at nickase-broken forks and *Brca1* Δ11 mutants revealed wild type levels of nickase-induced STGC ^38,44–46^. Taken together, current evidence suggests that DNA end resection at stalled or broken forks is less tightly regulated than at a replication-independent DSB, and that BRCA1 is largely dispensable for this process. The avid engagement of DNA end resection at stalled or broken forks may also explain why classical nonhomologous end joining (cNHEJ) does not compete with HR in nickase-induced broken fork repair or in stalled fork repair^1,38,47,48^.

Tandem duplications are common structural variants in the cancer genome, but the mechanisms that form them remain poorly understood ^6,40^. The ability to recapitulate *BRCA1*-linked Group 1 TD formation in the Tus/*Ter* system provides an opportunity to decipher mechanisms underlying this class of TDs. Our previous work led us to propose that Group 1 TDs form at a Tus/*Ter* RFB in cells lacking BRCA1 by a fork stalling/replication restart mechanism, resulting in re-replication of the TD segment, with completion of the TD by means of an end joining step ^1,6,7^ (**Supplemental Figure S7**). Our initial findings implicating the DNA end resection mediator CtIP as a Group 1 TD suppressor suggested that DNA end resection might play a role in Group 1 TD suppression^7^. In the current study, we approached this hypothesis by studying Tus/*Ter*-induced TD formation in different *Brca1* allelic backgrounds, in order to relate BRCA1-mediated Group 1 TD suppression to the known functions of BRCA1 in DSB repair. We find that, in contrast to *Brca1* Δ11 mutants, *Brca1* CC mutants are competent for suppression of Tus/*Ter*-induced TDs. This result further implicates the DNA end resection function of *Brca1* in Group 1 TD suppression. Described differently, Group 1 (∼10kb) TD formation is greatly accelerated when BRCA1-mediated DNA end resection capabilities are attenuated. We tested the predictive power of the Tus/*Ter* TD assay in a mouse model of *Brca*-linked mammary tumorigenesis ^25^.

Consistent with the predictions of the Tus/*Ter* system, we find that *Brca1* p.L1363P mutant-driven mammary tumors, although defective for HR, do not display a Group 1 TD phenotype. In the same experiment, confirming our previous work, *Brca1* null mammary tumors displayed a robust Group 1 TD phenotype^8^. In this regard, *Brca1* p.L1363P mutant cancers more closely resemble *Brca2* mutant cancers, which do not register as TDP ^8^. The Tus/*Ter* system accurately predicted these separation-of-function phenotypes of *Brca1* p.L1363P-driven cancers.

An important clue regarding the mechanism of Tus/*Ter*-induced TD formation came from our discovery of FANCM and BLM as Group 1 TD co-suppressors^7,28^. The synergistic induction of Tus/*Ter*-induced TDs in *Brca1* Δ11 cells lacking FANCM led us to study the genetic relationship between these two genes. We found that *Brca1* Δ11 mES cells are synthetic lethal/sick on a *Fancm* null or *Fancm* ATP hydrolysis-defective genetic background. If this relationship were to hold for *BRCA1*-linked cancers, small molecule inhibitors of FANCM ATPase function might be effective as therapeutics in *BRCA1*-linked cancer. These findings raised the possibility that Group 1 TD formation and sensitivity to FANCM loss might be mechanistically linked. Here, we tested this hypothesis by comparing the impact of *Fancm* deletion on the growth properties of *Brca1* Δ11 mutants, *Brca1* L1363P CC mutants and *Brca2* Δ27 mutant cells. By use of two different assays—colony plating efficiency and competitive growth assays—we confirm the synthetic sick phenotype of *Brca1* Δ11 *Fancm* null cells. In contrast, *Fancm* deletion in *Brca1* L1363P CC mutants was well-tolerated, with no defect in colony growth in comparison to *Brca1* L1363P CC mutants harboring WT *Fancm*. Indeed, *Brca1* L1363P *Fancm* null cells displayed a modest growth advantage over *Fancm* null cells that harbor WT *Brca1*. This counterintuitive finding is currently unexplained. However, interestingly, *BRCA1* p.L1363P-expressing cells were shown previously to exhibit increased frequencies of SSA^23^. Conceivably, enhanced SSA in *Brca1* L1363P cells might confer increased tolerance of *FANCM* loss in comparison to cells expressing WT *Brca1*. Overall, our results support the hypothesis that BRCA1-mediated DNA end resection, suppression of Group 1 TD formation and tolerance of *Fancm* loss are mechanistically linked (**Supplemental Figure S6C**). Analysis of additional *Brca1* mutant alleles would be helpful to explore this relationship further. Interestingly, we noted modest growth impairment of *Brca2* Δ27 cells following *Fancm* deletion, albeit less severe than the impact on *Brca1* Δ11 cells. *Brca2* Δ27 mutants disrupt a C-terminal RAD51 binding domain and are defective for classical HR, fork protection and gap suppression, but are competent for suppression of Tus/*Ter*-induced TDs^7,49,50^ (**Supplemental Figure S6B**). It will be interesting to study the impact of *Fancm* deletion in other *Brca2* allelic backgrounds.

If, as data here suggest, Group 1 TD formation and *Fancm* synthetic lethality are different consequences of defective BRCA1-mediated DNA end resection, how might these processes be linked? We suggest that FANCM and BRCA1 act as independent barriers to Group 1 TD formation: FANCM (in association with BLM^7,28^) acts by suppressing BIR-type replication restart at stalled forks; BRCA1, *via* its SSA function, provides a ‘failsafe’ mechanism for collapsing back to single copy any nascent Group 1 TDs that escape the action of FANCM/BLM (**Supplemental Figure S7**). SSA—a DSB repair function dependent on BRCA1-mediated DNA end resection but not on RAD51 loading—is traditionally considered an error-prone DSB repair pathway, since it can mediate genomic deletions between homologous repeats^1^. However, in the context of a nascent Group 1 TD, SSA would perform a conservative (genome-stabilizing) role by preventing TD formation (**Supplemental Figure S7B**). When both BRCA1 and FANCM are inactivated, two important, independent checks on Group 1 TD formation are lost, unleashing intolerable levels of genomic instability at sites of fork stalling (**Supplemental Figure S6C**).

One manifestation of this heightened genomic instability is an enhanced Group 1 TD phenotype; the other is the loss of cell fitness i.e., synthetic lethality.

## Limitations of the Study

This study is limited to mouse ES cells and mouse mammary tumors. One study in somatic human cells has also identified a synthetic lethal relationship between *BRCA1* and *FANCM* mutations ^29^. However, it is not yet clear to what extent the synthetic lethal relationship between *BRCA1* and *FANCM* translates to the cancer setting. The development of therapeutics that specifically target FANCM would facilitate such studies and might ultimately prove effective in the treatment of *BRCA1*-linked cancers. We have limited this study to mutations affecting *Brca1* exon 11 and the CC domain. It will be instructive to study additional domain mutants of BRCA1 and its associated proteins. As regards the mechanism of Group 1 TD formation, the link between BRCA1-mediated DNA end resection and TD suppression established by our work is correlative; it is formally possible that some other as yet unidentified function of BRCA1, which maps to the same domains as those required for DNA end resection, is the actual mediator of Group 1 TD suppression. However, the balance of evidence supports a role for DNA end resection in Group 1 TD suppression. The mechanisms discussed here apply only to Group 1 TDs, which have much smaller span sizes than most cancer-related TDs. Larger Group 2 or Group 3 TDs, which dominate the TD spectrum of non-*BRCA1*-linked cancers, likely arise by fundamentally different mechanisms, as reflected in their distinct genetic dependencies^6,8^.

The naturally occurring triggers of Group 1 TD formation in the cancer genome remain unknown. Although the Tus/*Ter* fork stalling system has thus far made accurate predictions regarding TD formation in *BRCA1*-linked cancer, it is possible that lesions additional to fork stalling lesions contribute to Group 1 TD formation in cancer. Indeed, Huang et al. recently showed that Group 1 TDs can form at a site-specific nick, implying that replication fork breakage can also result in Group 1 TD formation^51^. This finding raises the question: which form of replication stress—fork stalling or fork breakage—accounts for the major fraction of Group 1 TDs in a developing *BRCA1*-linked cancer? It will be instructive to compare the genetic regulation of Tus/*Ter*-induced *vs*. nickase-induced Group 1 TD formation and to relate these findings to the genetics of *BRCA1*-linked cancer-associated Group 1 TDs.

## Acknowledgements

We gratefully acknowledge helpful discussions with other Scully lab members and the contribution of the Genome Technologies Service and Data Science at The Jackson Laboratory for expert assistance with the WGS work described herein. This work was supported by NCI grants R35CA263813 to R.S., P30CA034196 and R01CA255705 to ETL, R01CA138804, R01CA262227 and P01CA250957 to BX, grants from Oncode Institute, the Dutch Cancer Society (NKI 2015-7877) and the Lundbeck Foundation (R223-2016-8) to J.J and an American Cancer Society Postdoctoral Fellowship to N.M.N.

## Author Contributions

All authors participated in the design of experiments. N.M.N, A.M.G., F.M., D.N., N.A.W. and E.W. performed the experiments. B.X. provided PALB2 antibodies. E.W. and J.J. provided tumor tissue for analysis. F.M and E.T.L. performed whole genome sequencing and bioinformatic analysis of tumor genomes. N.M.N., A.M.G., F.M. and R.S. wrote the first draft of the manuscript and all authors participated in manuscript editing.

## Disclosure and competing interests statement

R.S. is a member of the Scientific Advisory Board of MoMa Therapeutics, Inc.

## Materials and Methods

### Molecular biology, siRNA and sgRNA oligos

6x*Ter* HR reporters were engineered as previously described Using conventional cloning methods^7,27^. Plasmids used for transfection were prepared using endotoxin-free maxiprep (QIAGEN, 12362) kit and have been described previously^27^. *Brca1* exon 12 (L1363P) targeting construct was generated using large double-strand DNA gBlocks (Integrated DNA Technologies). Oligomers and primers used for genotyping, RT-qPCR and sgRNA synthesis were purchased from Thermo-Fisher Scientific and GeneWiz (Azenta). siRNA SMARTpools were purchased from Horizon Scientific/Dharmacon. Cas9-sgRNA target sites in the mouse genome sequence were identified using the Heidelberg CCtop tool: https://cctop.cos.uni-heidelberg.de.

sgRNA *in vitro* synthesis was performed using the Engen sgRNA Synthesis Kit (New England Biolabs, E3322S) and purified using the Clean and Concentrate Kit (Zymo Research, R1017). sgRNA was electrophoresed on a denaturing gel (Novex 10% TBE-Urea; ThermoFisher Scientific, EC6875BOX), to assess quality prior to transfection.

### Cell culture and cell lines

mES cells are thawed onto plates coated with mouse embryonic fibroblast (MEF) feeders, maintained in ES medium (DMEM with 15% serum, 20mM HEPES pH 7.6, 0.1mM bME, 500U/mL rLIF, 1x MEM NEAA, 2mM L-glutamine, 100U/mL penicillin, 100ug/mL streptomycin, 0.001M sodium pyruvate) on gelatinized plates, and regularly tested for mycoplasma infection by Myco Alert assay (Lonza, LT07-318). *Brca1*^fl5-13/+^ cells were targeted with a 6x*Ter*-HR reporter as a single copy to the *Rosa26* locus using standard methods described previously^27^. All cell lines for analysis of *Brca1* separation-of-function mutant alleles were derived from a *Brca1*^fl11/hyg^ founder cell line carrying one *Brca1* allele in which most of the conventionally termed exon 11 (NCBI exon 10) is replaced by a hygromycin resistance cassette, and one functionally wild-type conditional *Brca1* allele in which exon 11 is flanked by *LoxP* elements ^33^. The founder cell line contains a single copy of the 6x*Ter*-HR reporter, targeted as a single copy to the *Rosa26* locus^27^.

### Generation of *Brca1* mutant cell lines

To delete floxed alleles of *Brca1*, cells were treated with adeno-Cre for 24 h and then plated on 60 mm dishes with MEF feeder layer without any selection. Clones were picked after 4-5 days and expanded after for another 5 days. Genomic DNA from clones was prepared using Qiagen Puregene buffers and protocol. The clones were then assessed to check for recombined *LoxP* sites using primers specific for the deleted or undeleted conditional allele (Table 2). To generate *Brca1* hemizygous cell lines, sgRNA against NCBI exon 2 and 23 (Table 1) were designed and synthesized as described above. Cas9-sgRNA RNP was pre-assembled *in vitro* in OptiMEM by mixing 3 µM Spy NLS Cas9 (New England Biolabs, M0646T) and 3 µM purified sgRNAs. Cells were transfected using Lipofectamine 2000 mixed with RNP complexes. 48 h after transfection cells were plated onto 60 mm dishes containing feeder MEFs without selection. Individual clones were picked for expansion between 4-5 days and then expanded on MEFs for another 5 days.

**Table 1.**
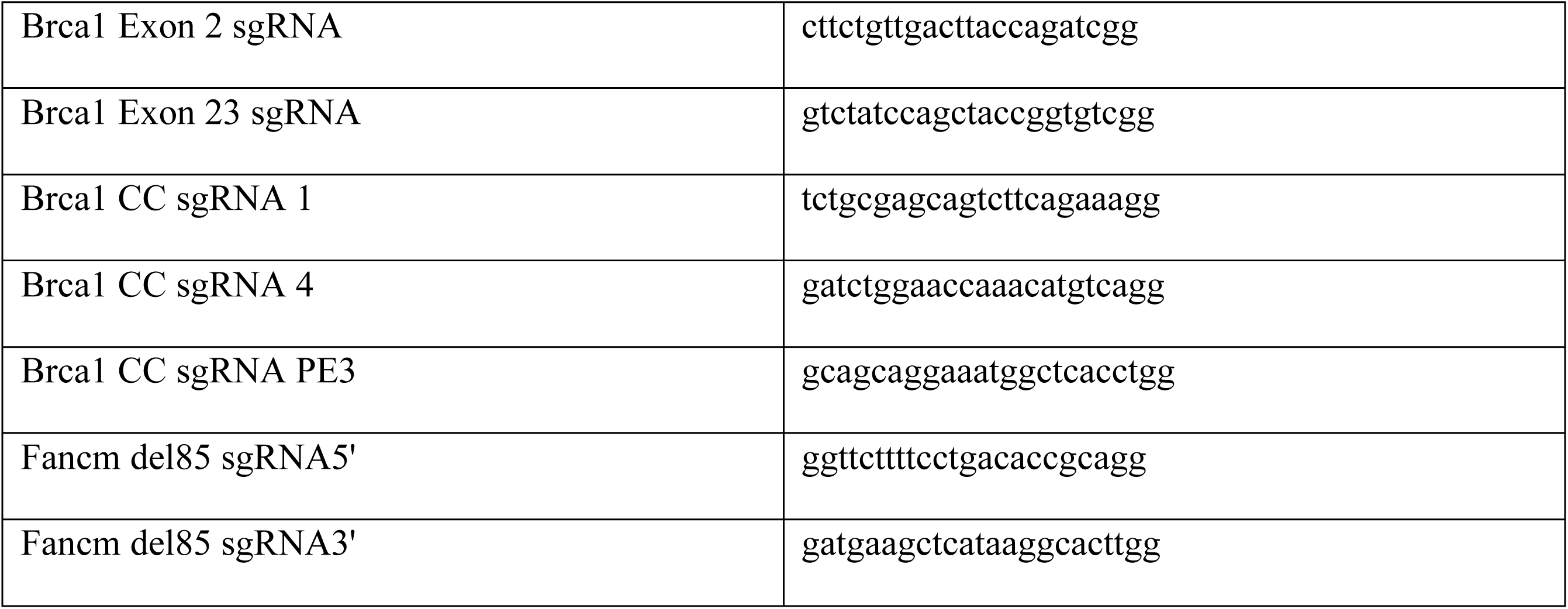
sgRNA oligomer sequence with PAM

**Table 2.**
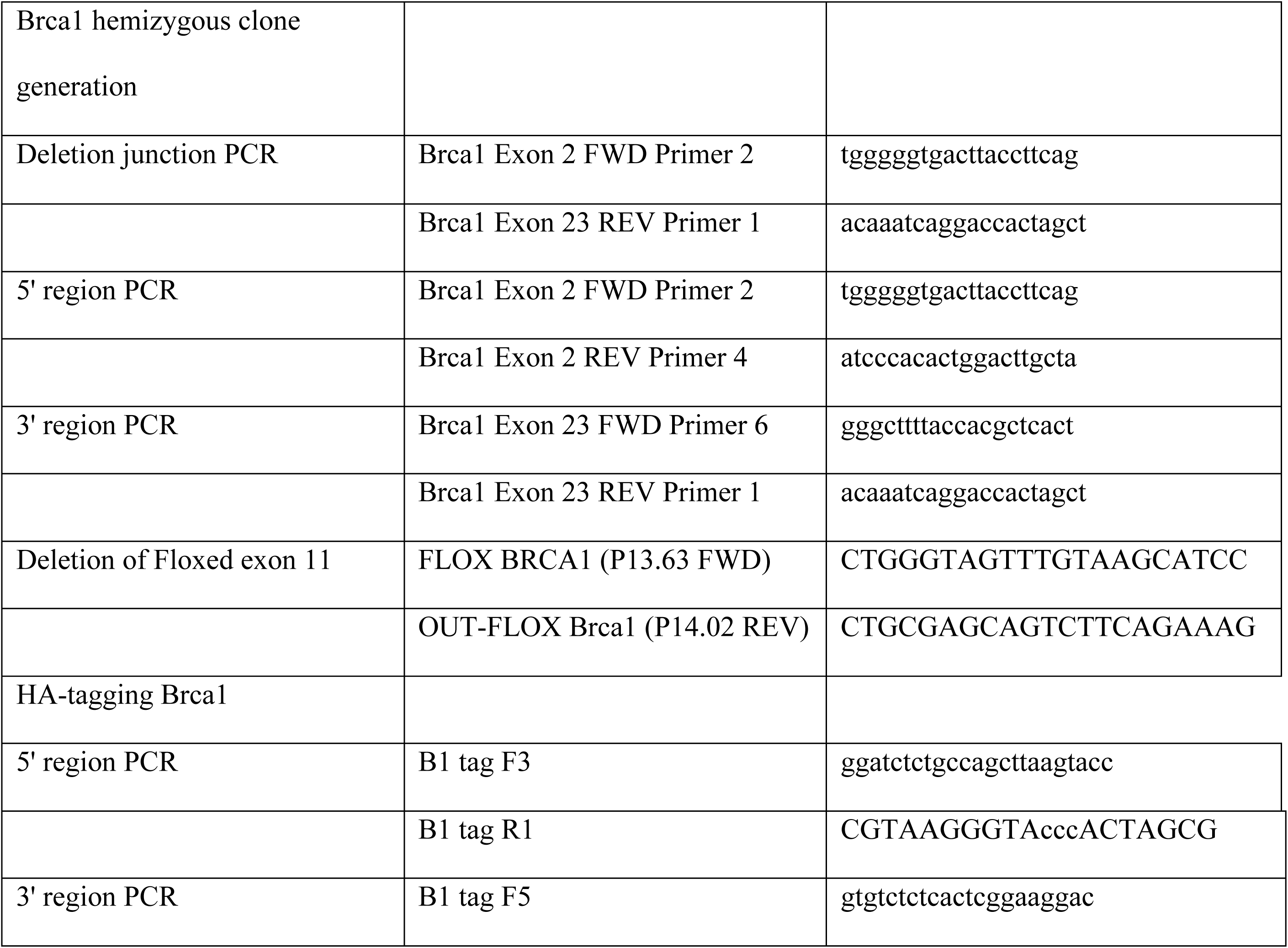
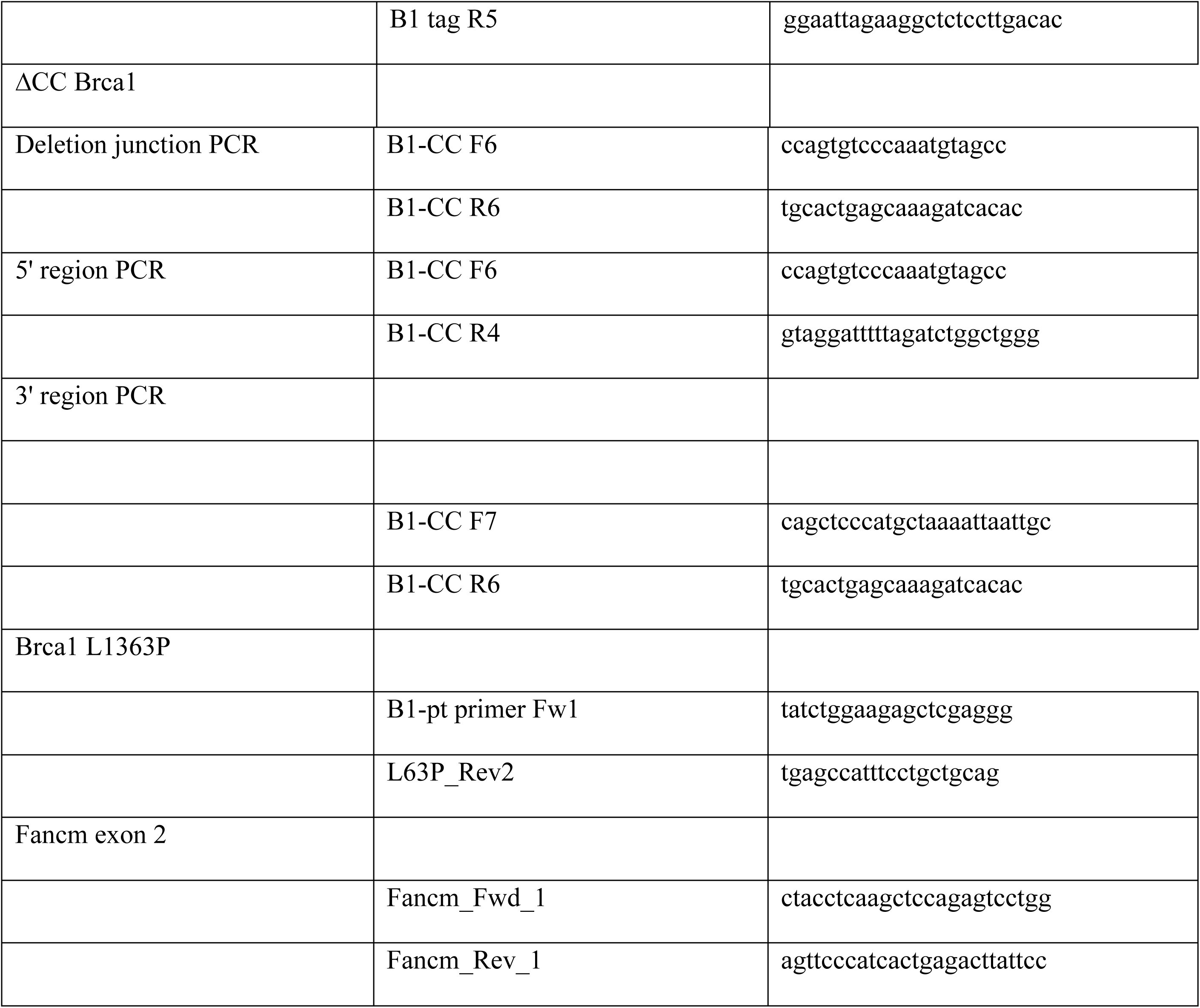
Primers used in the study

**Table 3.** Antibodies used in the study

|  |  |
| --- | --- |
| Anti-HA (for immunoblot) | Abcam ab9110 |
| Anti-HA (for immunofluorescence) | Abcam ab18181 |
| Anti-vinculin | CST #4650 |
| Anti-mouse BRCA1 exon11 | Scully lab |
| Anti-RPA (for ChIP) | Abcam ab10359 |
| Anti-mouse PALB2 | Xia lab |

*Brca1* hemizygous clones were identified by PCR and confirmed by sequencing using primers spanning the deleted *Brca1* allele (Table 2).

For in-frame *Brca1* HA-tagging, we generated a targeting construct which positions 8xHA tag sequences in-frame with the 3’ end of the *Brca1* open reading frame, followed by a SMASh degron and a neomycin resistance gene downstream of self-cleaving peptide T2A sequences. Cas9 RNP complexes targeting *Brca1* NCBI Exon 23 (Table 1) were co-transfected with a linearized targeting construct using Lipofectamine 2000. 48 h after transfection, cells were plated onto 60 mm dishes coated with a feeder layer of MEFs that express the neomycin resistance gene. 24 h after seeding, 4 µM G418 containing medium was added to the plates. 5-7 days later, G418-resistant colonies were isolated and expanded. Colonies were confirmed using PCR and Sanger sequencing using primers to detect successful integration of the HA-SMASh elements (Table 2). The clones were also assessed for full length *Brca1* gene product expression using immunoblotting against HA, as described in the Results section.

Domain mutations in the BRCA1 coiled coil were also generated using CRISPR/Cas9 gene editing. To generate ΔCC clones, sgRNAs targeting NCBI exon 11 and 12 (Table 1) were designed, synthesized *in vitro* and transfected as RNP complexes using Lipofectamine 2000. Clones were generated as mentioned above, were grown without selection and were screened by PCR followed by sequencing using primers for ΔCC Brca1 (Table 2).

*Brca1* p.L1363P point mutant clones were generated by transfecting a template containing specific mutations in a region of *Brca1* NCBI exon 12. For p.L1363P mutation, *ctg* was mutated to *ccg*, along with silent mutations in the PAM site of sgRNA used (Table 1). Primers *Brca1* L63P (Table 2) were used to identify clones through PCR and sequencing.

### Generation of mouse Fancm hemizygous and null clones

Characterized *Brca1*^HA/–^ (WT, L1363P and Δ11) and *Brca2*^Δ27/Δ27^ (deleted for exon 27) mutant clones were subjected to CRISPR/Cas9 gene editing to generate *Fancm+/–* and *Fancm–/–*clones. sgRNAs targeting exon 2 (Table 1) of the *Fancm* gene were designed to generate an 85 bp frame-shift deletion in the coding region of *Fancm*. Clones were grown without selection, isolated and expanded as described above. *Fancm* mutant clones and isogenic controls that retained WT *Fancm* were identified through PCR (Table 2) followed by Sanger sequencing the region of interest after TA cloning.

### Recombination assays

1.6 x 10^5^ mES cells were transfected in suspension with 0.5 μg empty vector or pcDNA3β-myc NLS-Tus, or pcDNA3β -myc NLS-I-SceI using Lipofectamine 2000 (Invitrogen) ^27,52^. Transfection efficiency was measured in parallel by transfecting 0.05 μg wild-type GFP expression vector and 0.45 μg control vector. For experiments with siRNAs, 0.3 µg of plasmid vectors were transfected along with 10 pmol siRNA for each of the target gene or along with 10 pmol of luciferase siRNA. 72 h after transfection, 3× 10^5^–6× 10^5^ total events were scored for each sample and GFP+RFP–, GFP+RFP+ and GFP–RFP+ frequencies were analyzed through flow cytometry using a Beckman Coulter CytoFlex, in duplicates. Reported repair frequencies were corrected for background events and for transfection efficiency (50–85%).

### Colony formation assays

To study the effect of PARPi on different *Brca1* mutants, approximately 800 cells were seeded onto 6 well tissue culture plates after coating them with 0.1% gelatin. 24 h after seeding the cells, medium was replaced and the plates were replenished with PARPi containing medium. 6 days after plating the cells, colonies were fixed using Carnoy’s fixative (75% methanol and 25% glacial acetic acid) and stained with crystal violet, the colonies were counted using a Celigo imaging cytometer (Revvity). For plating assays of *Fancm*^−/−^ clones, an equal number (400) of cells were plated on to 6 well cell culture dishes and allowed to grow for 6 days, after which the colonies were processed as described earlier.

### Competitive growth assays

0.04 x 10^6^ GFP^+^ *Brca1*^fl-HA/–^*;Fancm^−/−^*cells were mixed with 0.36 x10^6^ GFP^−^ *Brca* mutant *Fancm^−/−^* cells. Approximately, 0.02x10^6^ cells were seeded onto a 6 well plate for growth over two weeks and the GFP^+^ fractions were quantified by flow cytometry on the days shown in **Figure 7**.

### Immunoprecipitation

500 µL of NETN300 buffer with protease inhibitors was added onto a 60-70% confluent 100 mm cell culture dish and incubated on ice for 30 minutes. The lysate was then scraped and collected in a microfuge tube and spun down at 14K RPM in a refrigerated microcentrifuge at 4°C for 20 minutes. The supernatant was collected for immunoprecipitation. Approximately 600 µg of protein was added to 25 µL of anti-HA beads (Pierce) and rotated on an end-to-end rotator for 16-20 h at 4°C. The beads were then washed twice with lysis buffer, followed by one wash with 1xTBST (Tween20) and a final wash with ultrapure nuclease free water. The eluate was obtained by incubating the beads with 60 µL of 1x non-denaturing SDS loading buffer at 95 °C for 5 minutes. Samples were electrophoresed on a SDS-TAE 3-8% gradient gel overnight and the protein was transferred onto a nitrocellulose membrane for analysis by immunoblotting.

### Western blotting

RIPA buffer was added to cell pellet and incubated on ice for 30 minutes, following which the lysate was sonicated and spun down at 14K RPM in a refrigerated microcentrifuge at 4°C for 20 minutes. Protein was estimated using Thermo Rapid BCA Gold kit. Approximately, 30 µg of protein was used for electrophoresis in a TAE 3-8% gradient gel for 16-20 h overnight at 50 V in a cooled chamber. Protein was transferred onto a nitrocellulose membrane using wet transfer in a transfer buffer containing 20% methanol. Following transfer, the membrane was blocked in 5% skimmed milk for anti-HA and anti-vinculin immunoblots, whereas 5% BSA was used for anti-PALB2 immunoblot. After incubating the membranes with appropriate secondary antibody, Pico PLUS Chemiluminescent Substrate was used to visualize the immunoblot signals using a BioRad ChemiDoc MP.

### Nucleofection of CRISPR-Cas9 RNP

Equal volumes of 100 uM crRNA (Biosynthesis, IDT) and 100 uM tracrRNA (IDT) were mixed and incubated on a 95 C heat block for 3 mins, then cooled on a benchtop for 5 minutes. Dialysis buffer (20mM HEPES pH 7.5, 500mM KCl, 20% glycero), Cas9 (IDT, 10 ug/uL), Cas9 (IDT, 10 ug/uL), and annealed crRNA/tracrRNA were mixed to form RNP and allowed to sit at room temperature for at least twenty minutes. P3 solution with supplement (Lonza) were freshly mixed and used for nucleofection of trypsinized cells. 2 μL of 100 μM electroporation enhancer (IDT) was added to each nucleofection reaction. Approximately 12 x 10^6^ cells were used for each nucleofection.

### Chromatin-immunoprecipitation (ChIP)

Nucleofected cells were collected via cell scraping in plain DMEM. Cells were fixed with 1% formaldehyde for 10 minutes and gently agitated by rotation at room temperature and then quenched with 750 µL of 2M Glycine for 4 minutes with rotation at room temperature. Cells were spun at 1200 x *g* for 3 minutes at 4 C, washed with PBS, spun and flash frozen and stored at -80 C. Protein A magnetic beads (50 μL per sample) were washed with 0.5% BSA three times, then incubated with RPA antibody (abcam 10359, 5 μL per sample). The antibody was allowed to bind to the magnetic beads at room temperature with rotation for 1 to 3 hours. After incubation with antibody, beads were washed three times with 0.5% BSA in PBS. Cell pellets were removed from -80 °C and resuspended in lysis buffer (50 mM HEPES, 140 mM NaCl, 1 mM EDTA, 10% glycerol, 0.5% Igepal CA-630, 0.25% Triton-X; with pH adjusted to 7.5 using KOH and with 1x protease inhibitors immediately before use (ThermoFisher, 78429) for 10 minutes at 4 °C with rotation. The lysate was pelleted at 2000x *g* for 3 minutes at 4 °C, then resuspended in nuclear lysis buffer (10 mM Tris-HCl pH 8, 200 mM NaCl, 1 M EDTA and 0.5 mM EGTA; with pH adjusted to 8 using HCl and with 1x protease inhibitors added before use) and allowed to rotate for 5 minutes at 4 °C. The nuclear lysate was pelleted at 2000 x *g* for 3 minutes at 4 °C, then resuspended in sonication buffer (10 mM Tris-HCl pH 8, 100 mM NaCl, 1 mM EDTA, 0.5 mM EGTA, 0.1% Na-Deoxycholate, 0.5% N-lauroylsarcosine; with pH adjusted to 8.0 using HCl, and adding 1x protease inhibitor right before use) and sonicated using (Qsonica Q125)) with 30 seconds of pulse, 30 seconds off, 55% amplitude for 6 minutes at 4 °C. Chromatin was spun at 20000 x*g* at 4 °C. The supernatant was added to a fresh 5 ml lo-bind tube and 1% final volume of TrionX-100 and antibody bound beads were added. The final volume was increased to 3 mL using the sonication buffer. The mix was allowed to rotate for 8-20 hours at 4 °C.

After 8-20 hours of rotation, beads were collected and washed 6 times with RIPA solution (50 mM HEPES, 500 mM LiCl, 1 mM EDTA, 1 % Igepal CA-630, 0.7 % Na-Deoxycholate; pH adjusted to 7.5 using KOH), then once with TBS wash buffer (20 mM Tris-HCl pH 7.5, 150 mM NaCl) and resuspended in elution buffer (50 mM Tris-HCl pH 8, 10 mM EDTA, 1% SDS).

Beads and elution buffer were incubated for at least 8 hours with shaking at 65 °C for reverse crosslinking. RNaseA was added to the beads and incubated for 15 minutes at 37 °C, followed by Proteinase K at 55 C for 1 hour. Beads were separated from the eluate using magnetic separation and the DNA was purified using MinElute column purification.

The ChIP libraries were prepared according to the NEB Ultra II DNA Library Prep Kit (NEB, #E7645). End-repair and A-tailing was performed, followed by adaptor ligation (NEB 7600), and 13 cycles of PCR amplification. Samples were then pooled and DNA was quantified using a Qubit (ThermoFisher) and sequenced on an Illumina NextSeq 1000 machine using paired-end 2x60 bp read length.

### ChIP-Seq data analysis

ChIP-Seq data was demultiplexed using the bcl2fastq software and the resulting fastq files were aligned to a mouse genome (mm10) containing the HR reporter using bowtie2. Alignments were converted to binary, filtered by quality and sorted using samtools. Enrichment bigwig files were obtained using deeptools by subtracting the signal from the undamaged sample (No Cas9) to the damaged one (window size of 50 bp). Final plots (shown in **Figure 4B**) were generated using the PyGenomeTracks software. ChIP-Seq enrichments (**Figure 4C**) were computed using a custom python code that subtracts the number of normalized (RPM) reads in the damaged and undamaged sample in a window of 30 kb centered at the cut site. Analysis code is available on GitHub (https://github.com/AlbertoMarinG/BRCA1_SoF). Raw WGS data for ChIP-seq experiments are on the Sequence Read Archive (SRA: https://www.ncbi.nlm.nih.gov/sra), BioProject: PRJNA1378990; BioSamples: SAMN53860886.

### Mouse mammary tumor models and WGS

Mouse mammary tumors were generated in *K14-Cre;Trp53*^fl2-10/fl2-10^*;Brca1*^fl5-13/fl5-13^ (B1P), *K14-Cre;Trp53*^fl2-10/fl2-10^*;Brca2*^fl11/fl11^ (B2P) and *K14-cre;Trp53*^fl2-10/fl2-10^*;Brca1*^L1363P/fl5-13^ (Br1L1363P) female mice as described previously^8,25,32,39^. Genomic libraries were derived from 15 mouse tumor tissues and their matched mouse spleen tissues (normal controls), by shearing 200ng of genomic DNA using the R230 ultrasonicator (Covaris), followed by size selection using Ampure XP Beads. DNA fragments were then treated with end-repair, A-tailing, and ligation of Illumina unique adapters (Illumina) using the KAPA Hyper Prep Kit (Illumina) (KAPA Biosystems/ Roche), after which 5 cycles of PCR amplification were performed. Library quantification was performed using real-time qPCR (Thermo Fisher) before sequencing on a Novaseq X Plus Seq (Illumina) platform to generate 151-bp-long paired end reads, aiming for 30X coverage.

Sequencing reads were aligned to the mouse *GRCm39* reference genome using the Burrows-Wheeler Aligner^53^. For each cancer genome, we calculated the sequencing coverage across the *Trp53* and *Brca1* loci, extending each region by 50% of its length upstream and downstream, using non-overlapping 100-bp bins. Coverage values were then normalized across samples, using the *edgeR::calcNormFactors* function in R. A rolling average of the normalized coverage was computed for each sample, using windows of 25 and 50 consecutive 100-bp bins for *Trp53* and *Brca1*, respectively, as represented in **Supplemental Figure S5B**. We then averaged the values corresponding to the windows mapping to the delta region as well as the left and right flanking regions, for either gene, separately, and computed the delta allele frequency (DeltaFr) as 1-(coverage in the flanking windows/ coverage in the delta windows).

Structural variant calls were generated using four different tools (Manta^54^, Delly^55^, LUMPY^56^ and SvABA^57^, and high-confidence events were selected when called by at least two tools and by requiring split-read support. Matched normal DNA samples were used as controls to remove germline variants. A TDP score was computed for each tumor sample based on the number and chromosomal distribution of its somatic tandem duplications (TDs), as previously described^5^.

Single-nucleotide variants were called by MuTect2^58^, Strelka2^59^ and Lancet^60^. High confidence calls were defined as those that are either called by at least two variant callers or called by one caller and seen in the Lancet validation calls. Putative germline variants were removed by only keeping mutations unique to each cancer genome. In addition to the 15 tumor genomes sequenced as part of this study, we also applied the same pipeline to reanalyze 15 previously published mouse breast cancer genomes from the following genotypes: *K14-cre;Trp53*^fl2-10/fl2-10^ (KP, n = 3), *WAP-Cre;Trp53*^fl2-10/fl2-10^ (WP, n = 3), *K14-Cre;Trp53*^fl2-10/fl2-10^*;Brca1*^fl5-13/fl5-13^ (KB1P, n=3), *WAP-Cre;Trp53*^fl2-10/fl2-10^*;Brca1*^fl5-13/fl5-13^ (WB1P, n = 3), *K14-Cre;Trp53*^fl2-10/fl2-10^*;Brca2*^fl11/fl11^ (KB2P, n = 3)^5,8^.

Single base substitution (SBS) signatures were extracted with the *YAPSA* R package^61^ using the original 22 high confidence COSMIC signatures described by Alexandrov et al. as the mutational process matrix^62^.

WGS data relative to the new mouse breast cancers sequenced as part of this study (i.e. 15 pairs of spleen and tumor DNA samples) are available from the Sequence Read Archive database (www.ncbi.nlm.nih.gov/sra), SRA: PRJNA1393414. Previously published mouse cancer WGS datasets are also available from SRA: PRJNA430898.

---

**Figure S1.**
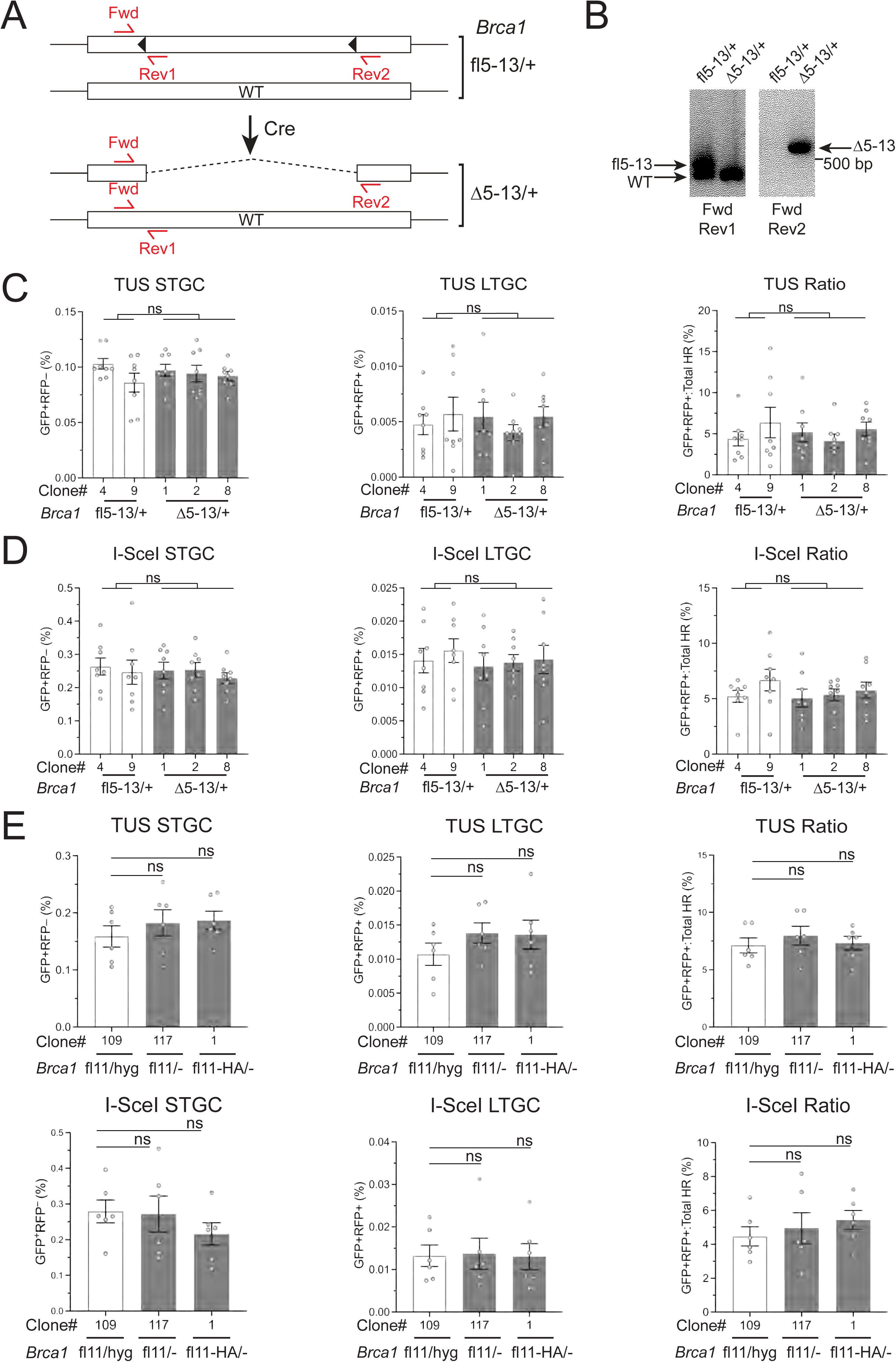
**Quantitation of HR in *Brca1* hemizygous mES cells**. **A**. Schematic of clones with one floxed *Brca1* allele, *Brca1*^fl5-13^ and Cre-mediated deletion product *Brca1*^Δ5-13^. Forward (Fwd) and reverse (Rev) primers used to detect deletion of the floxed region are shown. **B.** PCR genotyping of *Brca1*^fl5-13/+^ and *Brca1*^Δ5-13/+^ clones. **C-D.** HR repair product frequencies in three independent *Brca1*^Δ5-13/+^ clones and two independent *Brca1*^fl5-13/+^ isogenic clones. Data for Tus/*Ter*-induced (C) and I-Scel-induced (D) repair is shown. Data shows mean and SEM of eight independent experiments (n=8). Statistical analysis: RM-one way ANOVA with Geisser-Greenhouse correction: ns: not significant. **E.** Tus/*Ter*-induced (upper panel) and I-Scel-induced (lower panel) HR frequencies in parental *Brca1*^fl11/hyg^, *Brca1*^fl11/–^ and *Brca1*^fl11-HA/–^ are shown as mean and SEM of six independent experiments (n=6). Statistical analysis: Unpaired *t*-test with Welch’s correction. ns: not significant.

**Figure S2.**
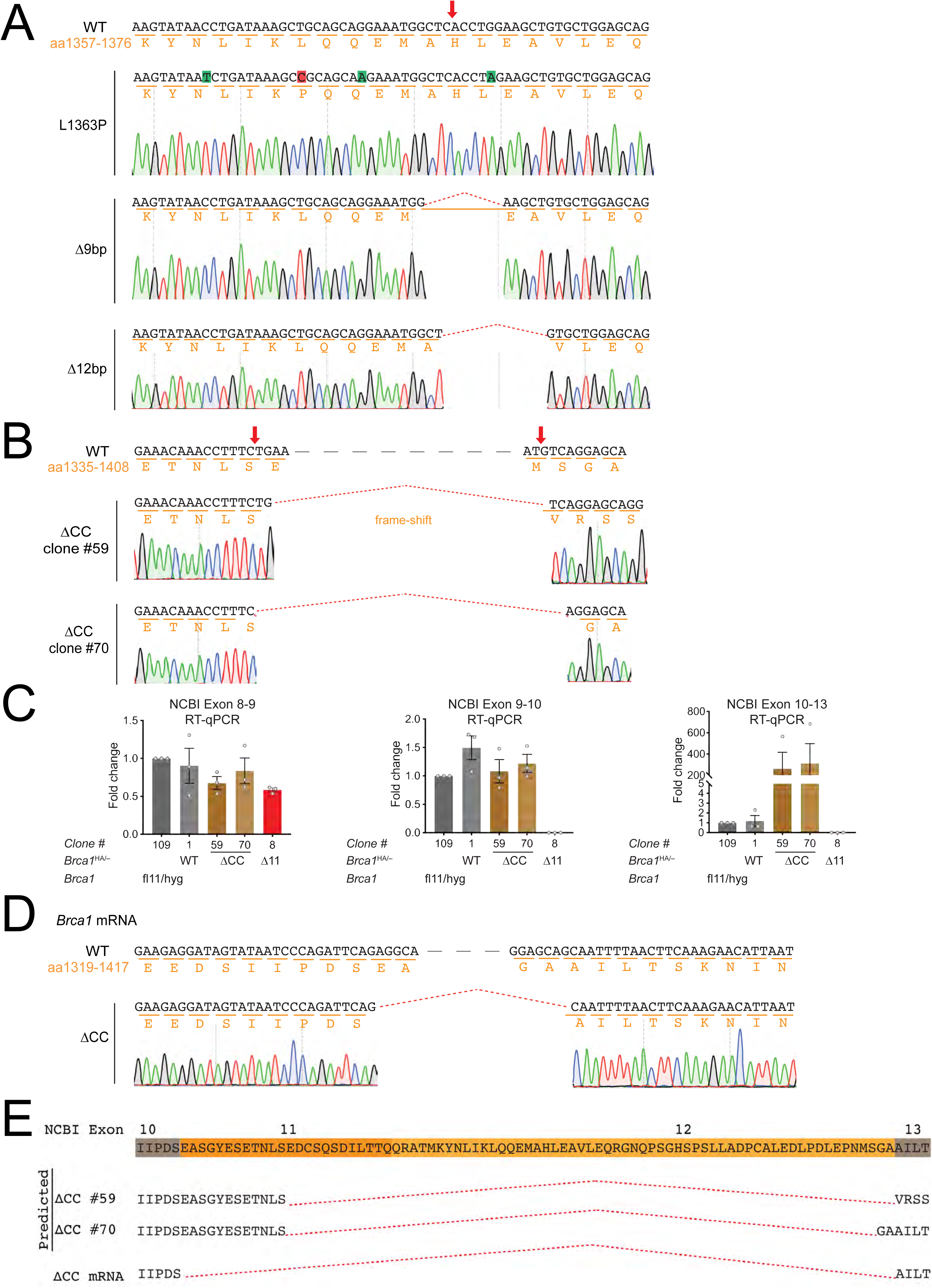
Sequence analysis of *Brca1* CC mutant clones and exon skipping in *Brca1*^ΔCC-HA/–^cells. *Brca1* allele abbreviations as in **Figure 2A**. Nucleotide sequence is shown in black and amino acids in orange. **A**. Nucleotide sequence of *Brca1* NCBI exon 12 encoding amino acids 1357-1376. Red arrow shows the predicted site of Cas9-induced DSB for generation of CC mutants. Red dotted line denotes deletion of nucleotides. Sequences of *Brca1* p.L1363P, *Brca1* CCΔ9bp and *Brca1* CCΔ12bp alleles are shown. L to P mutation in *Brca1* p.L1363P mutants is highlighted in red. Engineered silent mutations in potential sgRNA PAM sites are highlighted in green. **B**. *Brca1* NCBI exon 11 and 12 nucleotide sequence encoding amino acids 1335 to 1408. ΔCC clone #59 has a frame-shift mutation, whereas clone #70 is in-frame. **C.** RT-qPCR results showing amplification across three different exon-exon junctions (normalized to parental *Brca1*^fl11/hyg^ clone #109). Note absence of NCBI exon 10-11 amplification in Δ11 mutant clone and over-representation of NCBI exon 10-13 amplicon in ΔCC clones, as a result of exon skipping. **D**. Sequence of WT *Brca1* cDNA and *Brca1* ΔCC cDNA encompassing NCBI exons 10 to 13. Black dashed line represents continuation of sequence. Red dotted line represents deletion of nucleotides. ΔCC clones #59 and #70 revealed identical cDNA sequences, consistent with skipping of NCBI exons 11 and 12. **E**. BRCA1 amino acid sequence region encoded by 3’ end of *Brca1* NCBI exon 10 to 5’ end of NCBI exon 13. Amino acids encoded by NCBI exons 10 and 13 are highlighted in grey. Amino acids encoded by NCBI exons 11 and 12 are highlighted in orange and yellow, respectively. Predicted amino acid sequence based on sequencing of *Brca1* gDNA is shown for each ΔCC clone. Actual amino acid sequence based on *Brca1* ΔCC cDNA sequencing shows in-frame deletion resulting from skipping of NCBI exons 11-12.

**Figure S3.**
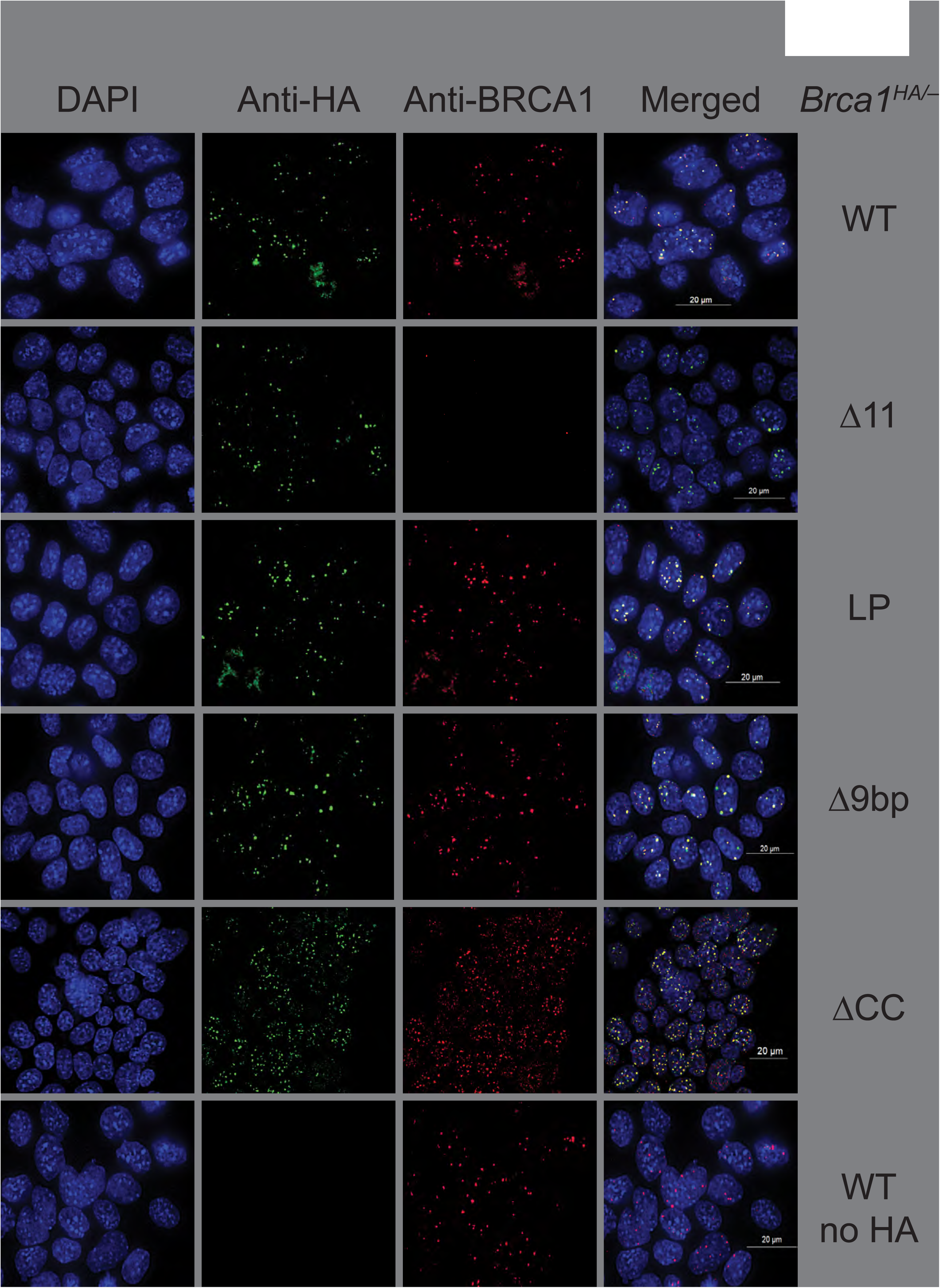
**Nuclear localization of BRCA1-8xHA mutants**. *Brca1* allele abbreviations as in **Figure 2A**. Immunofluorescence immunostaining of *Brca1*^HA/–^ mES cells. Alexa fluor 488 (green) was used to stain anti-HA signal, while Alexa fluor 594 (red) was used to stain for antibody raised against BRCA1 mouse Exon 11 (NCBI exon 10) gene product. Nucleus is counterstained using DAPI. Bottom row: WT *Brca1*^fl11/hyg^ mES cells (no HA signal). Note absence of red signal in *Brca1*^Δ11-HA/–^ cells and absence of green signal in *Brca1*^fl11/hyg^ cells. Images acquired using 60X objective. See STAR Methods for details.

**Figure S4.**
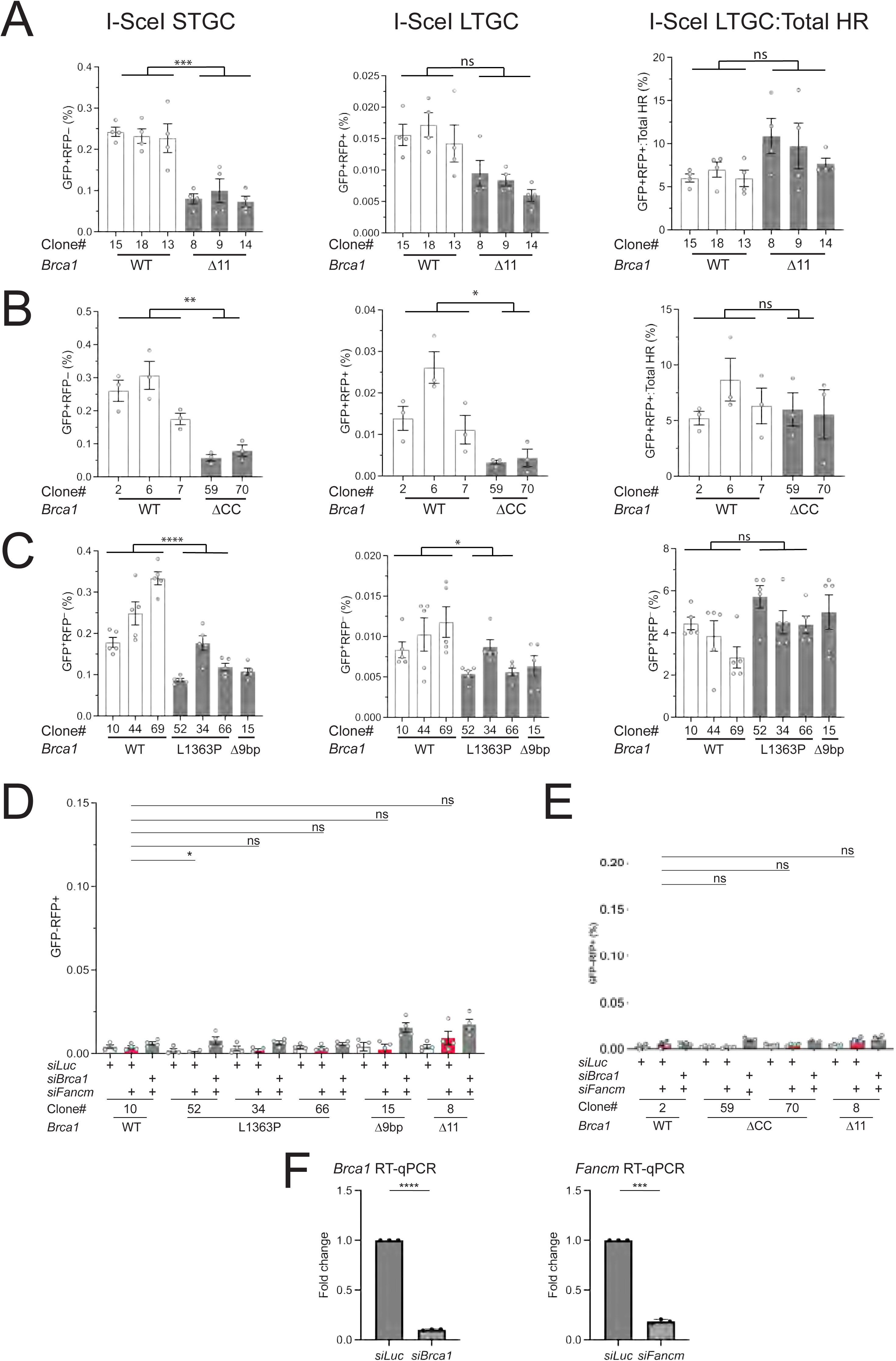
**DSB repair functions of *Brca1* CC mutants**. **A-C.** DSB-induced HR was quantified in *Brca1* mutants *vs*. isogenic WT clones. *Brca1* allele abbreviations as in **Figure 2A**. Data shows mean and SEM of n independent experiments. **A.** *Brca1*^Δ11-HA/–^ cells (n=4). **B.** *Brca1*^ΔCC-HA/–^ cells (n=3). **C.** *Brca1*^L1363P-HA/–^ and *Brca1*^CCΔ9bp-HA/–^ cells (n=5). Statistical analysis: RM-one way ANOVA with Geisser-Greenhouse correction: ns: not significant, * p <0.05, ** p <0.005, *** p < 0.0005, **** p <0.0001. **D** and **E**. I-SceI-induced GFP^–^RFP^+^ products in *Brca1* mutants. *Brca1* mutant cell lines were transfected with *I-SceI* plasmid together with 10 pmol of siRNAs against *Luciferase, Brca1* and *Fancm*. **D**. *Brca1*^L1363P-HA/–^ and *Brca1*^CCΔ9bp-HA/–^ mutants compared with their isogenic WT *Brca1*^fl11-HA/–^ clone and a *Brca1*^Δ11-HA/–^ clone. **E.** *Brca1*^ΔCC-HA/–^clones compared with their isogenic WT *Brca1*^fl11-HA/–^ clone and a *Brca1*^Δ11-HA/–^ clone (n=4). Statistical analysis: Unpaired *t*-test with Welch’s correction. ns: not significant, p * < 0.05. **F.** RT-qPCR analysis of *Brca1* and *Fancm* mRNA depletion following siRNA-mediated transfection. Statistical analysis: Unpaired *t*-test with Welch’s correction. *** p < 0.0005, **** p <0.0001.

**Figure S5.**
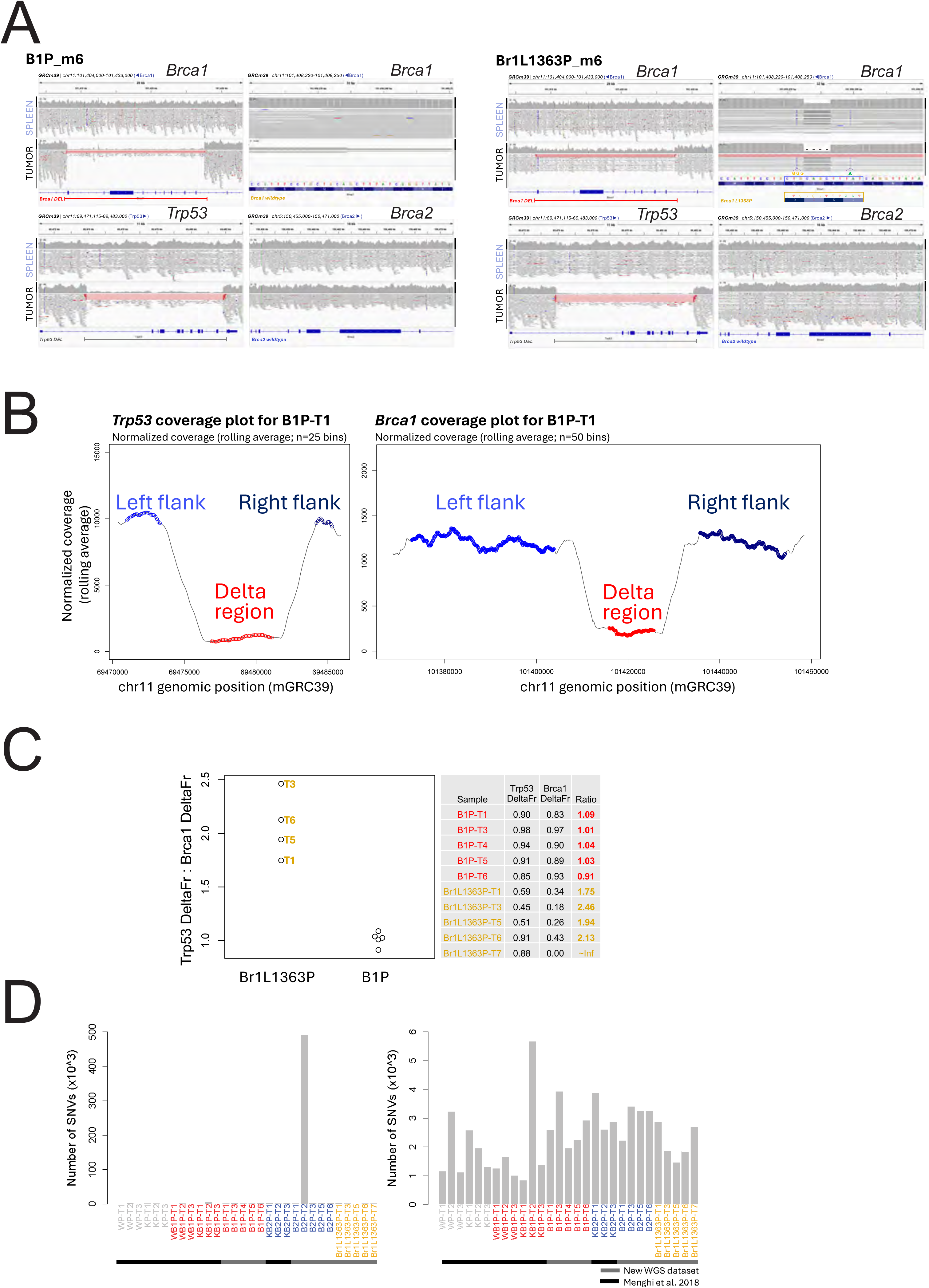
**Characterization of conditional *Brca* mutant tumors in *K14-Cre;****Trp53*^fl2-10/fl2-10^ **mice. A**. Integrative Genomic Viewer (IGV) screenshots of WGS reads mapping to the targeted genomic regions in matched spleen and mammary tumor tissues from individual animals representative of *K14-Cre;Trp53*^fl2-10/fl2-10^*;Brca1*^fl5-13/fl5-13^ (B1P) and *K14-cre;Trp53*^fl2-10/fl2-10^*;Brca1*^L1363P/fl5-13^ (Br1L1363P) mouse models examined in this study. For each screenshot, coverage and alignment tracks are highlighted by black and gray vertical bars, respectively. Bars and associated annotations below the gene structure track identify the exact genomic coordinates for the detected deletions. For the *Brca1* p.L1363P mutation, annotations have been added to clarify the nucleotide and amino-acid sequences corresponding to both the wild type and mutant *Brca1* alleles. Note that since the *Brca1* gene is encoded on the negative strand, the reported nucleotide sequence corresponds to the reverse and complement of the actual coding sequence. **B**. Examples of coverage plots for *Trp53* and *Brca1* targeted regions, showing marked drop in coverage for the genomic segments corresponding to the Cre-induced deletions (Delta region in red) compared to the left and right flanking regions (light and dark blue, respectively), which was quantified as described in the Methods section. **C.** The *Trp53*-DeltaFr:*Brca1*-DeltaFr ratio was computed to confirm the genotypes of the Br1L1363P and B1P tumors. Computed as 1– the average ratio between the coverage of the flanking region and that of the delta region, DeltaFr here indicates the extent of the loss of coverage associated with the Cre-induced deletion of the target regions. A DeltaFr = 1 corresponds to a decrease in coverage down to 0; a DeltaFr = 0 corresponds to equal coverage between flanking and delta regions. As expected for tumors homozygous for the *Trp53*^Δ2-10^ allele and heterozygous for the *Brca1*^Δ5-13^ allele, Br1L1363P tumors have a *Trp53*-DeltaFr:*Brca1*-DeltaFr ratio of about 2. In contrast, in the B1P tumors, which are homozygous for both the *Trp53*^Δ2-10^ and *Brca1*^Δ5-13^ alleles, the *Trp53*-DeltaFr:*Brca1*-DeltaFr ratio = ∼1. The only exception to this pattern is the Br1L1363P tumor 7, which contained an outsized deletion of *Brca1* that included the defined flanking regions and hence could not be plotted since the *Trp53*-DeltaFr:*Brca1*-DeltaFr ratio is infinite. **D.** Numbers of somatic single nucleotide variants (SNV) per tumor sample across the 15 mouse cancer genomes previously published^1^ and the new mouse cancer genomes sequenced as part of this study, including (left) or excluding (right) the B2P-T2 outlier sample.

**Figure S6.**
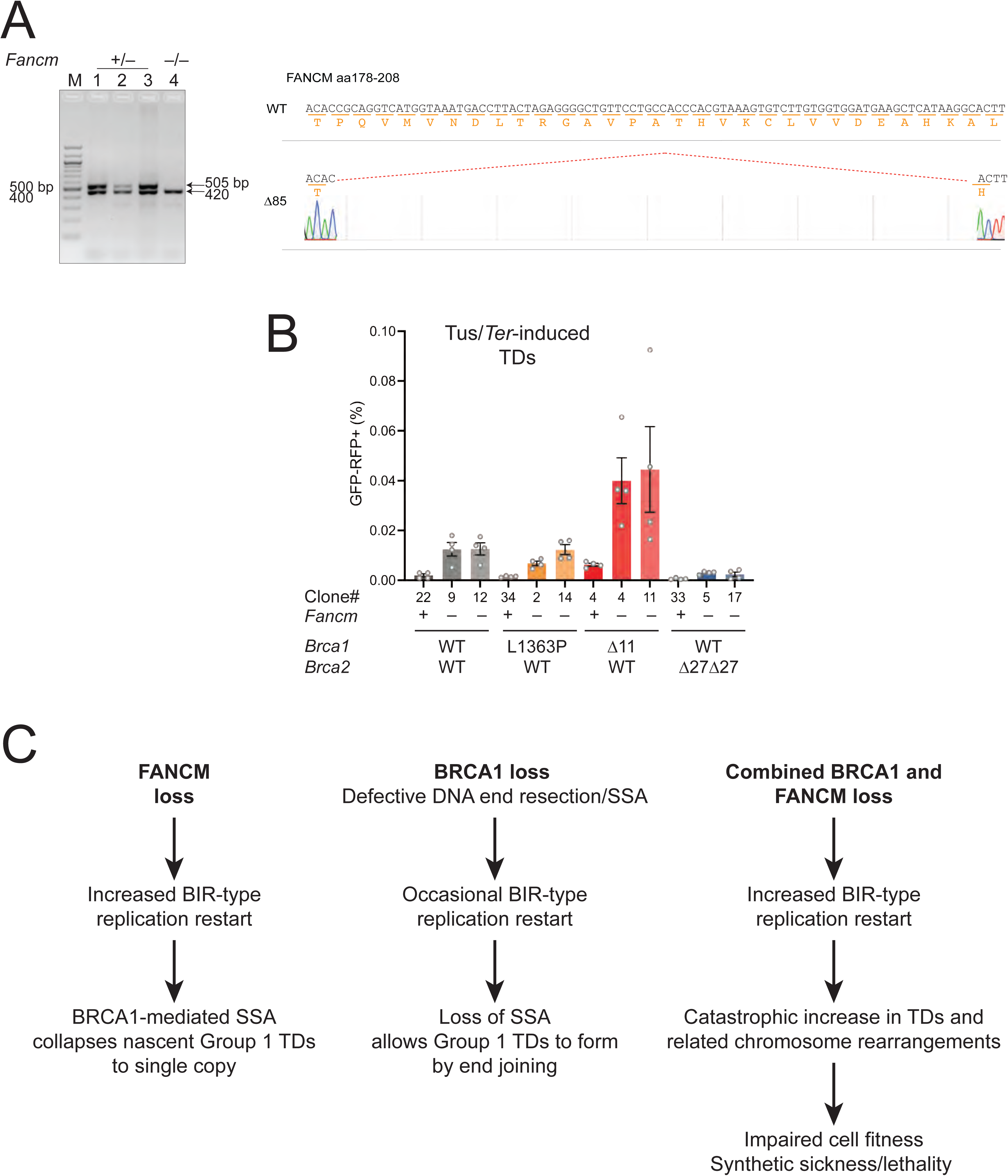
Genetic interactions between *Brca* mutant alleles and *Fancm* in mES cells. **A.** Genotyping of *Fancm*^+/–^ and *Fancm*^−/−^ clones. Left panel: PCR analysis of three *Fancm*^+/–^ clones and one *Fancm*^−/−^ clone. Right: sequence of *Fancm* Δ85 allele. This allele was shown previously to be functionally null^2^. **B.** Effect of siRNA-mediated *Fancm* depletion on Tus/*Ter*-induced TD formation in different *Fancm*^−/−^ *Brca* mutant clones. Note induction of TDs in *Fancm*^−/−^ *Brca1*^Δ11-HA/–^ clones and basal (WT) TD levels in other genotypes. **C.** Hypothesis linking BRCA1-mediated DNA end resection to *Fancm* synthetic lethality. Left: FANCM loss alone permits BIR-type replication restart at sites of stalled replication. BRCA1-mediated SSA collapses nascent Group TDs to single copy. Middle: Loss of BRCA1 alone abrogates SSA and permits occasional BIR-restarted replication forks to resolve as Group 1 TDs. Right: combined loss of BRCA1 and FANCM massively increases frequency of Group 1 TD formation at sites of replication stalling. The increase in Group 1 TDs and related chromosome rearrangement outcomes impairs cell fitness, resulting in synthetic sickness/lethality in the double mutant background.

**Figure S7.**
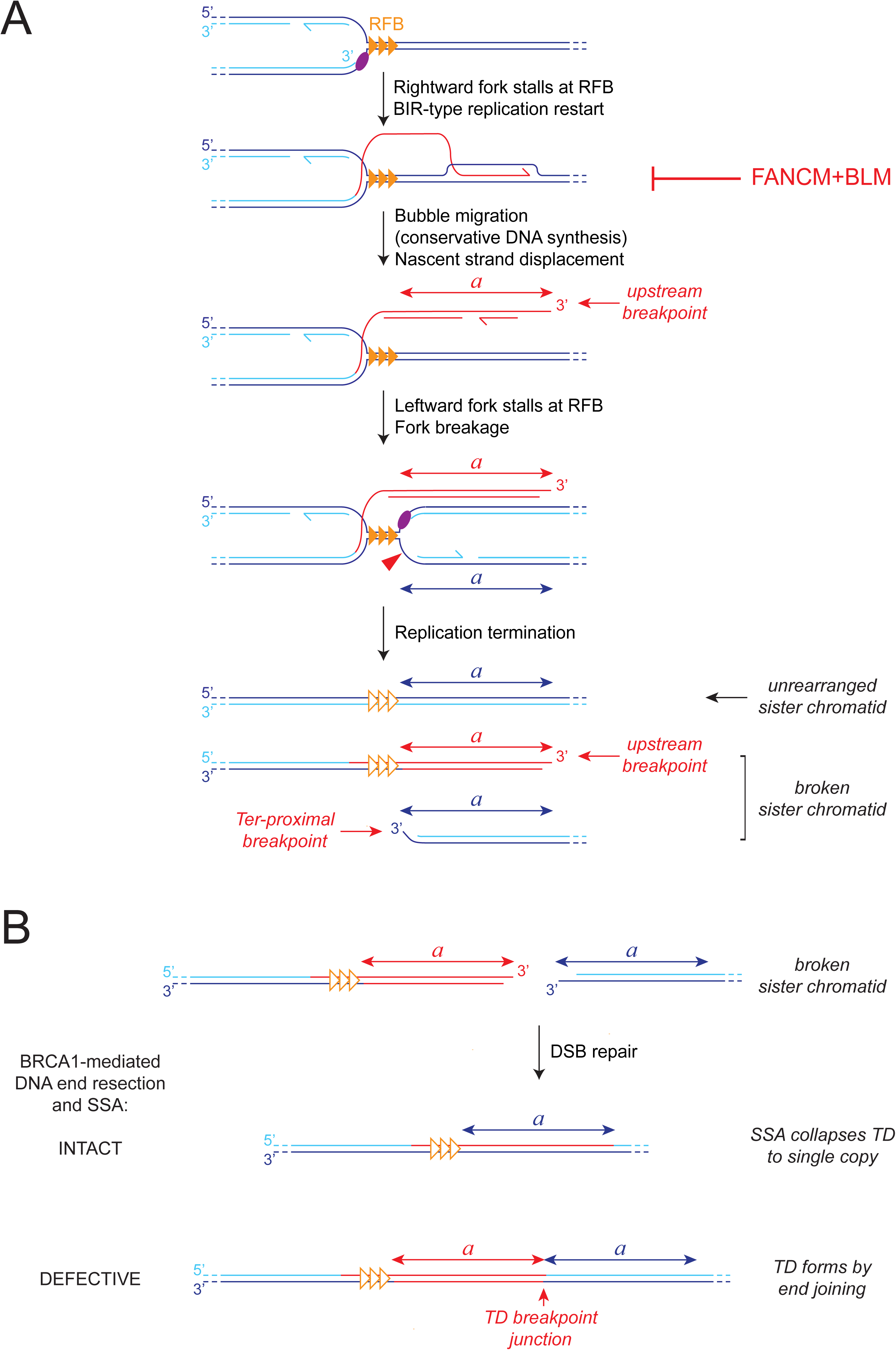
Proposed mechanisms of action of FANCM and BRCA1 in suppression of Tus/*Ter*-induced Group 1 TDs. **A.** Interaction of converging replication forks with Tus/*Ter* replication fork barrier (RFB) sets up the Group 1 TD intermediate. The Tus/*Ter*-induced TD breakpoint junction is formed from one breakpoint in close proximity to the *Ter* array (the ‘*Ter*-proximal’ breakpoint) and one ‘upstream’ breakpoint several kilobases from the *Ter* array^3^. We infer that for one fork (the rightward fork shown here), replication extends beyond the Tus/*Ter* RFB by an aberrant replication restart mechanism, while the second fork (here, the leftward fork) stalls and is processed at the Tus/*Ter* RFB site. We propose that the rightward fork stalls at the RFB and is restarted as a migrating bubble (i.e., BIR-type conservative DNA synthesis; red). This process could be *RAD51*-dependent or -independent^2,4^. FANCM, in partnership with BLM, suppresses this form of replication restart by disassembling migrating bubble BIR intermediates. Eventual termination of BIR generates a displaced nascent strand that spans the potentially duplicated segment (*a*). The 3’ end of the displaced nascent strand forms the potential upstream breakpoint of the TD. The leftward fork stalls at the RFB and either undergoes fork breakage as shown or fork reversal (not shown). The resulting solitary DNA end forms the potential *Ter*-proximal breakpoint. Completion of DNA replication through the RFB generates one intact sister chromatid and one broken sister chromatid that contains two copies of the TD segment *a*, flanking the break site. **B.** BRCA1 acts on the broken sister chromatid to suppress Group 1 TD formation. If BRCA1-mediated DNA end resection and SSA functions are intact, the two copies of the duplicated segment *a* flanking the break site are collapsed to single copy by BRCA1-mediated SSA. If BRCA1-mediated SSA is disabled, end joining opportunistically repairs the break, generating a Group 1 TD. The breakpoint junction is formed by end joining between the upstream breakpoint and the *Ter*-proximal breakpoint.

## References

1. Scully, R., Panday, A., Elango, R. & Willis, N.A. DNA double-strand break repair-pathway choice in somatic mammalian cells. Nat Rev Mol Cell Biol 20, 698–714 (2019).

2. Chen, C.C., Feng, W., Lim, P.X., Kass, E.M. & Jasin, M. Homology-Directed Repair and the Role of BRCA1, BRCA2, and Related Proteins in Genome Integrity and Cancer. Annu Rev Cancer Biol 2, 313–336 (2018).

3. Nik-Zainal, S. et al. Landscape of somatic mutations in 560 breast cancer whole-genome sequences. Nature 534, 47–54 (2016).

4. Lord, C.J. & Ashworth, A. PARP inhibitors: Synthetic lethality in the clinic. Science 355, 1152–1158 (2017).

5. Menghi, F. et al. The tandem duplicator phenotype as a distinct genomic configuration in cancer. Proc Natl Acad Sci U S A 113, E2373–82 (2016).

6. Scully, R., Glodzik, D., Menghi, F., Liu, E.T. & Zhang, C.Z. Mechanisms of tandem duplication in the cancer genome. DNA Repair (Amst) 145, 103802 (2025).

7. Willis, N.A. et al. Mechanism of tandem duplication formation in BRCA1-mutant cells. Nature 551, 590–595 (2017).

8. Menghi, F. et al. The Tandem Duplicator Phenotype Is a Prevalent Genome-Wide Cancer Configuration Driven by Distinct Gene Mutations. Cancer Cell 34, 197–210 e5 (2018).

9. Zhao, W., Wiese, C., Kwon, Y., Hromas, R. & Sung, P. The BRCA Tumor Suppressor Network in Chromosome Damage Repair by Homologous Recombination. Annu Rev Biochem 88, 221–245 (2019).

10. Cruz-Garcia, A., Lopez-Saavedra, A. & Huertas, P. BRCA1 accelerates CtIP-mediated DNA-end resection. Cell Rep 9, 451–9 (2014).

11. Yu, X., Wu, L.C., Bowcock, A.M., Aronheim, A. & Baer, R. The C-terminal (BRCT) domains of BRCA1 interact in vivo with CtIP, a protein implicated in the CtBP pathway of transcriptional repression. J Biol Chem 273, 25388–92 (1998).

12. Cejka, P. & Symington, L.S. DNA End Resection: Mechanism and Control. Annu Rev Genet 55, 285–307 (2021).

13. Ceppi, I. et al. Mechanism of BRCA1-BARD1 function in DNA end resection and DNA protection. Nature (2024).

14. Salunkhe, S. et al. Promotion of DNA end resection by BRCA1-BARD1 in homologous recombination. Nature (2024).

15. Jensen, R.B., Carreira, A. & Kowalczykowski, S.C. Purified human BRCA2 stimulates RAD51-mediated recombination. Nature 467, 678–83 (2010).

16. Thorslund, T. et al. The breast cancer tumor suppressor BRCA2 promotes the specific targeting of RAD51 to single-stranded DNA. Nat Struct Mol Biol 17, 1263–5 (2010).

17. Bell, J.C., Dombrowski, C.C., Plank, J.L., Jensen, R.B. & Kowalczykowski, S.C. BRCA2 chaperones RAD51 to single molecules of RPA-coated ssDNA. Proc Natl Acad Sci U S A 120, e2221971120 (2023).

18. Shahid, T. et al. Structure and mechanism of action of the BRCA2 breast cancer tumor suppressor. Nat Struct Mol Biol 21, 962–968 (2014).

19. Dray, E. et al. Enhancement of RAD51 recombinase activity by the tumor suppressor PALB2. Nat Struct Mol Biol 17, 1255–9 (2010).

20. Xia, B. et al. Control of BRCA2 cellular and clinical functions by a nuclear partner, PALB2. Mol Cell 22, 719–29 (2006).

21. Sy, S.M., Huen, M.S. & Chen, J. PALB2 is an integral component of the BRCA complex required for homologous recombination repair. Proc Natl Acad Sci U S A 106, 7155–60 (2009).

22. Zhang, F. et al. PALB2 links BRCA1 and BRCA2 in the DNA-damage response. Curr Biol 19, 524–9 (2009).

23. Anantha, R.W. et al. Functional and mutational landscapes of BRCA1 for homology-directed repair and therapy resistance. Elife 6(2017).

24. Zhao, W. et al. BRCA1-BARD1 promotes RAD51-mediated homologous DNA pairing. Nature (2017).

25. Pulver, E.M. et al. A BRCA1 Coiled-Coil Domain Variant Disrupting PALB2 Interaction Promotes the Development of Mammary Tumors and Confers a Targetable Defect in Homologous Recombination Repair. Cancer Res 81, 6171–6182 (2021).

26. Nacson, J. et al. BRCA1 Mutational Complementation Induces Synthetic Viability. Mol Cell 78, 951–959.e6 (2020).

27. Willis, N.A. et al. BRCA1 controls homologous recombination at Tus/Ter-stalled mammalian replication forks. Nature 510, 556–9 (2014).

28. Panday, A. et al. FANCM regulates repair pathway choice at stalled replication forks. Mol Cell 81, 2428–2444.e6 (2021).

29. Adam, S. et al. The CIP2A-TOPBP1 axis safeguards chromosome stability and is a synthetic lethal target for BRCA-mutated cancer. Nat Cancer 2, 1357–1371 (2021).

30. Setton, J. et al. Long-molecule scars of backup DNA repair in BRCA1- and BRCA2-deficient cancers. Nature 621, 129–137 (2023).

31. Egli, D. et al. Inter-homologue repair in fertilized human eggs? Nature 560, E5–e7 (2018).

32. Liu, X. et al. Somatic loss of BRCA1 and p53 in mice induces mammary tumors with features of human BRCA1-mutated basal-like breast cancer. Proc Natl Acad Sci U S A 104, 12111–6 (2007).

33. Xu, X. et al. Conditional mutation of Brca1 in mammary epithelial cells results in blunted ductal morphogenesis and tumour formation. Nat Genet 22, 37–43 (1999).

34. Bunting, S.F. et al. 53BP1 inhibits homologous recombination in Brca1-deficient cells by blocking resection of DNA breaks. Cell 141, 243–54 (2010).

35. Chung, H.K. et al. Tunable and reversible drug control of protein production via a self-excising degron. Nat Chem Biol 11, 713–20 (2015).

36. Scully, R. et al. Location of BRCA1 in human breast and ovarian cancer cells. Science 272, 123–6 (1996).

37. Huber, L.J. et al. Impaired DNA damage response in cells expressing an exon 11-deleted murine Brca1 variant that localizes to nuclear foci. Mol Cell Biol 21, 4005–15. (2001).

38. Elango, R. et al. Two-ended recombination at a Flp-nickase-broken replication fork. Mol Cell 85, 78–90.e3 (2025).

39. Jonkers, J. et al. Synergistic tumor suppressor activity of BRCA2 and p53 in a conditional mouse model for breast cancer. Nat Genet 29, 418–25 (2001).

40. Li, Y. et al. Patterns of somatic structural variation in human cancer genomes. Nature 578, 112–121 (2020).

41. Morimatsu, M., Donoho, G. & Hasty, P. Cells deleted for Brca2 COOH terminus exhibit hypersensitivity to gamma-radiation and premature senescence. Cancer Res 58, 3441–7 (1998).

42. Farmer, H. et al. Targeting the DNA repair defect in BRCA mutant cells as a therapeutic strategy. Nature 434, 917–21 (2005).

43. Chandramouly, G. et al. BRCA1 and CtIP suppress long-tract gene conversion between sister chromatids. Nat Commun 4, 2404 (2013).

44. Pavani, R. et al. Structure and repair of replication-coupled DNA breaks. Science 385, eado3867 (2024).

45. Scully, R., Walter, J.C. & Nussenzweig, A. One-ended and two-ended breaks at nickase-broken replication forks. DNA Repair (Amst) 144, 103783 (2024).

46. Zong, D., Pavani, R. & Nussenzweig, A. New twist on BRCA1-mediated DNA recombination repair and tumor suppression. Trends Cell Biol (2025).

47. Willis, N.A., Panday, A., Duffey, E.E. & Scully, R. Rad51 recruitment and exclusion of non-homologous end joining during homologous recombination at a Tus/Ter mammalian replication fork barrier. PLoS Genet 14, e1007486 (2018).

48. Vriend, L.E. et al. Distinct genetic control of homologous recombination repair of Cas9-induced double-strand breaks, nicks and paired nicks. Nucleic Acids Res (2016).

49. Longo, M.A. et al. BRCA2 C-terminal clamp restructures RAD51 dimers to bind B-DNA for replication fork stability. Mol Cell 85, 2080–2096.e6 (2025).

50. Lim, P.X., Zaman, M., Feng, W. & Jasin, M. BRCA2 promotes genomic integrity and therapy resistance primarily through its role in homology-directed repair. Mol Cell 84, 447–462.e10 (2024).

51. Huang, Z.C., et al. Late steps of allelic break-induced replication suppress tandem duplication associated with BRCA1 deficiency. Nucleic Acids Res 53(2025).

52. Puget, N., Knowlton, M. & Scully, R. Molecular analysis of sister chromatid recombination in mammalian cells. DNA Repair (Amst) 4, 149–61 (2005).

53. Li, H. & Durbin, R. Fast and accurate short read alignment with Burrows-Wheeler transform. Bioinformatics 25, 1754–60 (2009).

54. Chen, X., et al. Manta: rapid detection of structural variants and indels for germline and cancer sequencing applications. Bioinformatics 32, 1220–2 (2016).

55. Rausch, T. et al. DELLY: structural variant discovery by integrated paired-end and split-read analysis. Bioinformatics 28, i333–i339 (2012).

56. Layer, R.M., Chiang, C., Quinlan, A.R. & Hall, I.M. LUMPY: a probabilistic framework for structural variant discovery. Genome Biol 15, R84 (2014).

57. Wala, J.A. et al. SvABA: genome-wide detection of structural variants and indels by local assembly. Genome Res 28, 581–591 (2018).

58. Benjamin, D. et al. Calling Somatic SNVs and Indels with Mutect2. bioRxiv (2019).

59. Kim, S. et al. Strelka2: fast and accurate calling of germline and somatic variants. Nat Methods 15, 591–594 (2018).

60. Narzisi, G. et al. Genome-wide somatic variant calling using localized colored de Bruijn graphs. Commun Biol 1, 20 (2018).

61. Hübschmann, D. et al. Analysis of mutational signatures with yet another package for signature analysis. Genes Chromosomes Cancer 60, 314–331 (2021).

62. Alexandrov, L.B. et al. Signatures of mutational processes in human cancer. Nature 500, 415–21 (2013).

## References

1 Menghi, F., Barthel, F.P., Yadav, V., Tang, M., Ji, B., Tang, Z., Carter, G.W., Ruan, Y., Scully, R., Verhaak, R.G.W., et al. (2018). The Tandem Duplicator Phenotype Is a Prevalent Genome-Wide Cancer Configuration Driven by Distinct Gene Mutations. Cancer Cell 34, 197–210 e195. 10.1016/j.ccell.2018.06.008.

2. Panday, A., Willis, N.A., Elango, R., Menghi, F., Duffey, E.E., Liu, E.T., and Scully, R. (2021). FANCM regulates repair pathway choice at stalled replication forks. Mol Cell 81, 2428–2444.e2426. 10.1016/j.molcel.2021.03.044.

3. Willis, N.A., Frock, R.L., Menghi, F., Duffey, E.E., Panday, A., Camacho, V., Hasty, E.P., Liu, E.T., Alt, F.W., and Scully, R. (2017). Mechanism of tandem duplication formation in BRCA1-mutant cells. Nature 551, 590–595. 10.1038/nature24477.

4. Scully, R., Elango, R., Panday, A., and Willis, N.A. (2021). Recombination and restart at blocked replication forks. Curr Opin Genet Dev 71, 154–162. 10.1016/j.gde.2021.08.003.

